# Further Exploration of the SAR of Imidazoquinolines; Identification of Potent C7 Substituted Imidazoquinolines

**DOI:** 10.1101/793232

**Authors:** Jordan R. Hunt, Peter A. Kleindl, K. Ryan Moulder, Thomas E. Prisinzano, M. Laird Forrest

## Abstract

Small molecule agonists of TLR7/8, such as imidazoquinolines, are validated agonists for the treatment of cancer and for use in vaccine adjuvants. Imidazoquinolines have been extensively modified to understand the structure-activity relationship (SAR) at the N1- and C2-positions resulting in the clinical drug imiquimod, resiquimod, and several other highly potent analogues. However, the SAR of the aryl ring has not been fully elucidated in the literature. This initial study examines the SAR of C7-substituted imidazoquinolines. These compounds not only demonstrated that TLR7/8 tolerate changes at the C7 position but can increase potency and change their cytokine profiles. The most notable TLR7/8 agonists developed from this study **5**, **8**, and **14** which are up to 4-fold and 2-fold more active than resiquimod for TLR8 and/or TLR7, respectively, and up to 100-fold more active than the FDA approved imiquimod for TLR7.

---

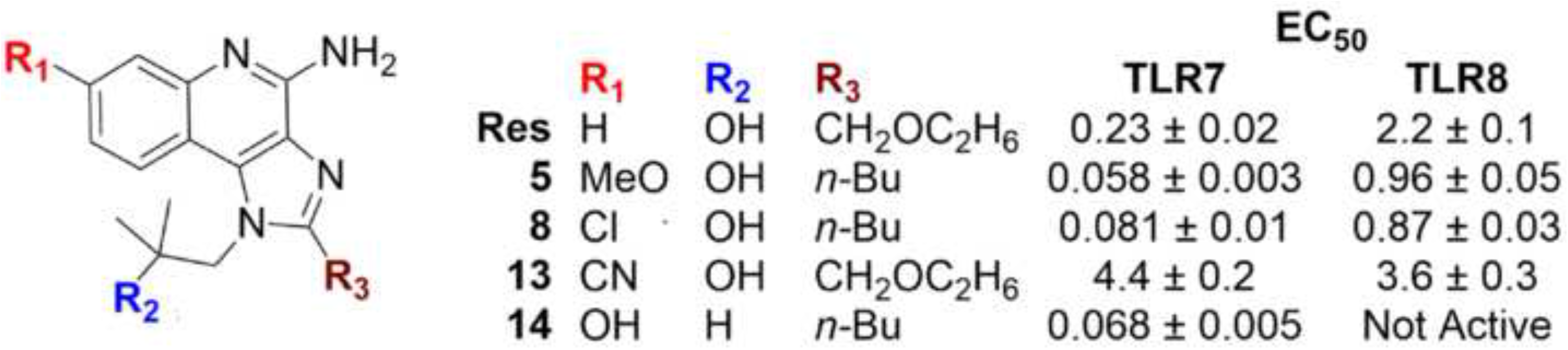

Toll-like Receptors (TLRs) are a validated target for developing new immunostimulatory drugs.^1,2^ TLRs are a part of the innate immune system, called pattern recognition receptors (PRRs), that help prime the adaptive immune system.^3^ PRRs detect pathogen-associated molecular patterns (PAMPs), which are conserved features common to many pathogenic microorganisms and viruses, but not expressed by the host organism (e.g. flagellin, lipopolysaccharides, bacterial DNA, and viral RNA).^4^ In humans, there are 10 TLRs (1-10), which can broadly be separated into endosomal (TLR3, 7, 8 and 9) and cell surface receptors (TLR1, 2, 4, 5, 6 and 10).^4^ The endosomal TLRs are natively activated by nucleic acid constructs not normally found or tightly controlled in the host, including double stranded RNA (TLR3), single stranded RNA (TLR7/8) and unmethylated CpG oligodeoxynucleotides (TLR9).^5^ TLR7/8 are expressed in a subset of immune system leukocytes, including monocytes, macrophages, natural killer (NK) cells, and B- and T-lymphocytes.^6^ TLR7/8 are canonical activators of the MyD88 pathway, which induces expression of pro-inflammatory type I interferons.^7^ Activation of Toll-like Receptor 7 and 8 (TLR7/8) is associated with a Th-1 biased (cell-mediated) immune response, as opposed to a Th-2 humoral (antibody) response.^8^ The cell-mediated immune response primarily helps eliminate intracellular pathogens, such as viruses, that may have limited exposure to humoral defenses.^9^ The cell-mediated immune response is also crucial in the host regulation of improper cell replication and development of cancers.^10^ For these reasons, small molecule agonists of TLR7/8 are highly sought after for the development of new vaccine adjuvants or antitumor agents.

Native ligands of TLR7/8 are guanosine and uridine-rich sequences common to viral single stranded RNA (ssRNA).^11^ In the case of TLR7, guanosine binds a first docking site on the receptor, which primes a second site to bind uridine with subsequent dimerization and activation.^12^ Binding of either guanosine or uridine alone is insufficient for activation. However, several small molecules have been described with sufficient affinity to activate TLR7/8 after binding of the first (guanosine) site alone, including imidazoquinolines, thiazoquinolines, benzoazepines, benzonaphthyridine, and guanine analogs.^13–16^ The most recognizable imidazoquinolines are imiquimod and resiquimod (**Figure 1**). Imiquimod is the only FDA-approved TLR7 agonist and is the active ingredient in the topical ointment Aldara^®^, used to treat skin conditions such as superficial basal cell carcinoma and actinic keratosis.^17^ Resiquimod is a dual TLR7/8 agonist that is in clinical trials for treatment of cutaneous T-cell lymphoma and melanoma.^18–20^

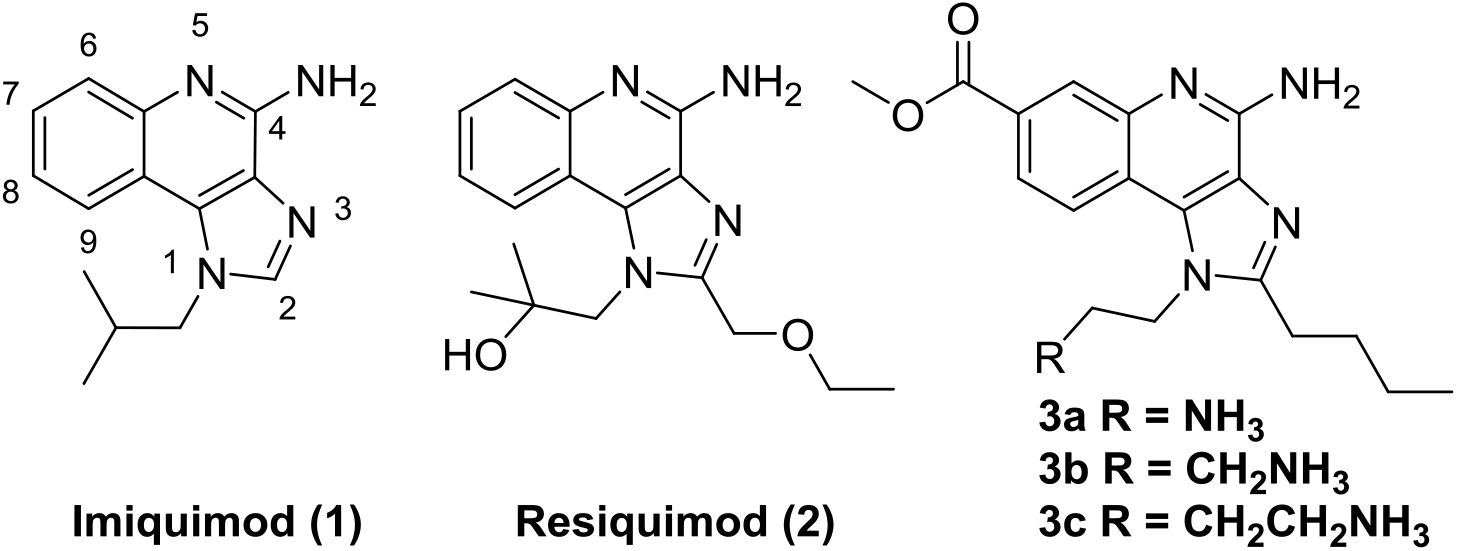

Since imidazoquinolines have been proven to be useful immunostimulants, significant effort has been put forth to develop the structure-activity relationship (SAR) and to help identify potent and effective imidazoquinolines suitable for vaccine adjuvants and cancer treatments.^21–25^ To date, the SAR investigations of imidazoquinolines have mostly focused on the N1 and C2 positions. The N1 position has been substituted with several different groups with differing results of selective TLR7, TLR8 or mixed TLR7/8 agonists.^22,24–26^ TLR7 tolerates many changes at N1 and C2 and are not required for activity, however, substitutions at these positions typically increase potency.^26,27^ TLR8 typically prefers N1 alkyl hydroxy substituents and either C2 *n*-butyl or *n*-pentyl with *n*-pentyl being more potent to produce mixed TLR7/8 agonists.^24^ The C4 amine is required for activity as changes to this position result in no activity.^26^ One position that has been relatively under-explored is the C7 position. Gerster et al. found that imidazoquinolines with a C7 methoxy, hydroxyl, and/or methyl group retained similar IFNα production when compared to **1**.^26^ Recently, Larson et al. showed that imidazoquinolines **3a-c** are TLR8 selective.^25^ In addition, the disclosure of the crystal structure of **2** bound to the TLR8 receptor provides further evidence that the C7 position might tolerate changes better than other aryl positions.^28^ To investigate this hypothesis, we synthesized various C7 substituted derivatives and evaluated their activities in TLR7/8 reporter cell lines and their cytokine induction in donor canine leukocytes. In this initial study, we explored the influence of electronic effects at the C7 position through the addition of electron withdrawing and electron donating groups (EWGs and EDGs).

Our initial synthetic targets for this investigation were aimed at C7 methoxy (**4** and **5**), chloro (**6-9**), nitrile (**10-13**), and hydroxyl (**14**) analogs as shown in Figure 2. The C7 methoxy group was selected due to its strong electron donating properties as well as its hydrogen bond accepting ability. The C7 hydroxy was chosen for its strong electron donating properties and its hydrogen bond accepting and donating abilities. The C7 chloro was picked for its lipophilic properties as well as its resonance into the aryl ring system. The C7 nitrile was selected for its strong electron withdrawing properties and polarizing nature. Also, in this study, the N1 substituent was changed from an isobutyl to the 2-hydroxy-2-methylpropyl while the C2 was changed from the *n*-butyl and ethoxymethyl. These N1 and C2 substitutions were mixed to observe if changes at C7 would change the known SAR of the N1 and C2 positions.

**Figure 2.**
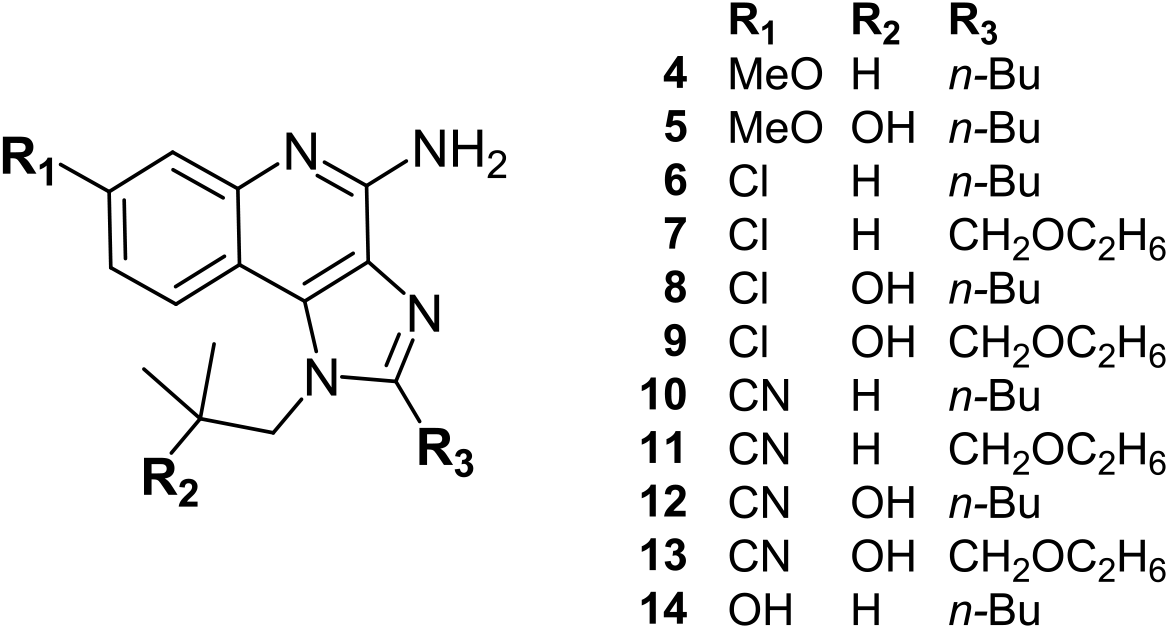
Target imidzoquinolines

**Scheme 1.**
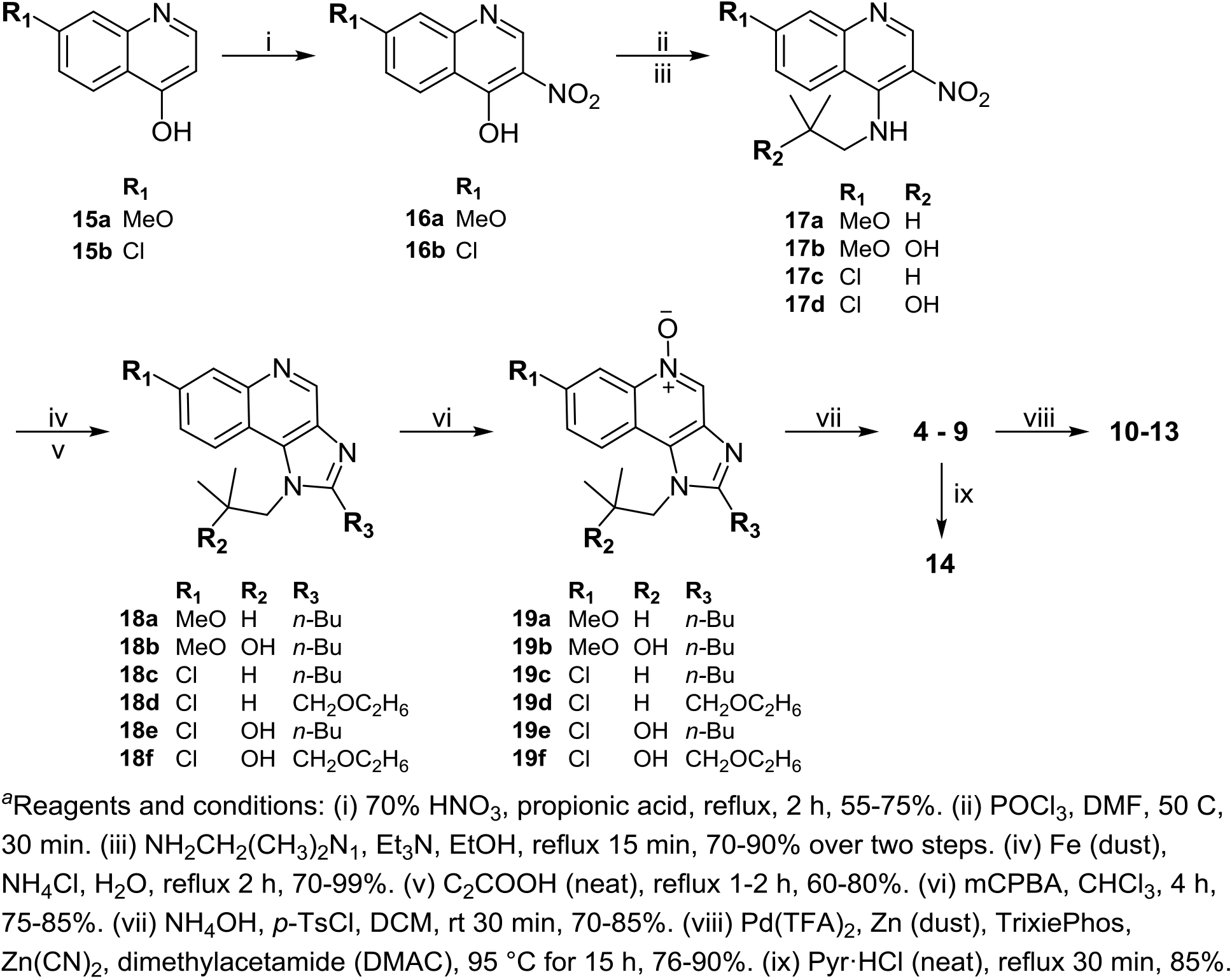
Synthesis of 7-substitued immidazoquinolines^*a*^

To obtain our targets, synthesis was started with the commercially available 7-methoxyquinolin-4-ol (**15a**) and 7-chloroquinolin-4-ol (**15b**) outlined in Scheme 1. **15a** and **15b** were subjected to 70% HNO_3_ in propionic acid to obtain the 3-nitroquinolinols (**16a,b**). Next, **16a** and **16b** were chlorinated using POCl_3_ and alkylated with the appropriate aminoalkyl chain to obtain **17a-d**. Reduction of the nitro group in **17a-d** using iron dust and ammonia chloride gave the diamines which were immediately used in the subsequent step to form the imidazoquinolines **18a-f** by condensation with the appropriate acid under refluxing conditions. *N*-oxides **19a-f** were made by oxidizing **18a-f** with *m*-CPBA. Subsequent treatment of **19a-f** with *p*-TsCl and ammonium hydroxide afforded compounds **4-9**. The overall yield for the 7-step sequence of reaction was approximately 30%. The C7 chloro analogues (**6-9**) were converted into the corresponding C7 nitriles (**10-13**) in yields of at least 76% using a mixture of Pd(TFA)2, rac-2-(Di-tert-butylphosphino)-1,1′-binaphthyl (TrixiePhos), Zn dust and Zn(CN)2.^29^ Finally, the demethylation of **4** to phenol **14** was accomplished in 85% yield using pyridinium hydrochloride.

The analogs (**4 – 14**) were screened for TLR7 and TLR8 activity *in vitro* using commercially available human embryonic kidney cells (HEK293) stably co-transfected to express either hTLR7 or hTLR8, along with a secreted embryonic alkaline phosphatase (SEAP) reporter gene induced by NF-κB (HEK-Blue^TM^, InvivoGen). TLR activation induces NF-κB production and subsequent SEAP secretion, which is measured using HEK-Detection media (InvivoGen) containing a colorimetric substrate for SEAP (UV-Vis, 637 nm). Transfected null cell lines lacking the hTLR7 or hTLR8 receptors but possessing the NF-κB induced SEAP reporter gene were used as a control to screen for non-specific activation of NF-κB (**Table 1**). Our target compounds were not evaluated at concentrations higher than 100 μM due to solubility issues.

**Table 1.** TLR7/8 Agonist Activities of Analogs **4-13**^*a*^

| Compound | EC<sub>50</sub> (μM) ± SD<sup>b</sup> |  | Cytokine (pg/mL) ± SD<sup>b</sup> |  |  |  |  |  |
| --- | --- | --- | --- | --- | --- | --- | --- | --- |
|  | TLR7 | TLR8 | TNFα | IL-12 | IL-6 | IL-10 | IL-2 | IFNγ |
| **1** | 6.8 ± 0.8 | - | 25 ± 3 | 1200 ± 140 | 300 ± 48 | 16 ± 2 | 85 ± 37 | 18 ± 6 |
| **2** | 0.25 ± 0.03 | 3.0 ± 0.3 | 190 ± 30 | 2800 ± 520 | 380 ± 80 | 21 ± 2 | 120 ± 12 | 26 ± 7 |
| **4** | 0.11 ± 0.01 | >100<sup>c</sup> | 49 ± 9 | 746 ± 130 | 500 ± 140 | 37 ± 13 | 30 ± 29 | 0 |
| **5** | 0.081 ± 0.009 | 1.7 ± 0.3 | 64 ± 12 | 2000 ± 160 | 290 ± 63 | 14 ± 2 | 82 ± 56 | 14.5 ± 6 |
| **6** | 0.3 ± 0.02 | >100 | 91 ± 20 | 760 ± 50 | 240 ± 67 | 14 ± 4 | 63 ± 44 | 7 ± 1 |
| **7** | 0.91 ± 0.09 | >100 | 114 ± 25 | 781 ± 40 | 280 ± 81 | 20 ± 1 | 50 ± 22 | 5 ± 5 |
| **8** | 0.084 ± 0.007 | 1.5 ± 0.2 | 22 ± 4 | 1700 ± 330 | 230 ± 44 | 7 ± 6 | 55 ± 30 | 13 ± 10 |
| **9** | 0.63 ± 0.03 | 4.9 ± 0.6 | 70 ± 8 | 1900 ± 130 | 320 ± 76 | 12 ± 3 | 76 ± 59 | 16 ± 2 |
| **10** | 0.99 ± 0.40 | >100 | 10 ± 1 | 920 ± 170 | 350 ± 95 | 37 ± 10 | 60 ± 41 | 13 ± 5 |
| **11** | 6.8 ± 0.7 | >100 | 7 ± 1 | 190 ± 62 | 71 ± 20 | 6 ± 2 | 57 ± 14 | 4 ± 3 |
| **12** | 0.79 ± 0.23 | 3.9 ± 0.2 | 44 ± 10 | 418 ± 72 | 290 ± 77 | 14 ± 3 | 45 ± 22 | 12 ± 4 |
| **13** | 3.5 ± 0.7 | 27 ± 6 | 80 ± 20 | 13 ± 3 | 214 ± 69 | 6 ± 3 | 46 ± 42 | 0.2 ± 0.4 |
| **14** | 0.073 ± 0.005 | >100 | 270 ± 70 | 102 ± 65 | 330 ± 92 | 17 ± 3 | 79 ± 19 | 9 ± 8 |
| **Vehicle** |  |  | 8 ± 2 | 18 ± 8 | 310 ± 90 | 33 ± 16 | 47 ± 20 | 0.4 ± 0.7 |

<sup>a</sup>Average EC<sub>50</sub> values were determined from three independent measurements at 10 or more concentrations, performed in quadruplicate using either hTLR7 or hTLR8 transfected HEK-Blue cells along with corresponding Null controls. Raw data and statistical methods available in SI. <sup>b</sup>Cytokine production was measured in triplicate at 20 μM, full graphs and statistical analysis available in SI. <sup>c</sup>>100 = no activity observed at 100 μM.

As expected, imiquimod (**1**) and resiquimod (**2**) were found to be agonists for TLR7 and TLR8 receptors with values consistent with previous reports.^23–26^ Generally, we found that (1) TLR7 is more tolerant to changes than TLR8; (2) tertiary alcohols at the N1 position tend to increase the TLR7 activity when compared to those without the tertiary alcohol; and (3) TLR8 activity requires the tertiary alcohol at N1.^23^ This is all consistent with previous investigations.^22–26^ We also observed that when C2 contains the ether substitution, TLR7/8 activity decreases when compared to the C2-butyl. This suggests that the changes made at C7 do not cause a change in the trend for N1 and C2 sites.

As others have found, TLR7/8 continues to tolerate C7 substitutions when the N1 and C2 are substituted appropriately.^24^ The most important finding is that there is a general trend for C7 substitutions. It has been observed in this study that EDGs are stronger activators of TLR7/8 than EWGs. For TLR7 the range was from no statistically significant difference to a 13-fold decrease. For TLR8 the range was from no statistically significant difference to a 9-fold decrease. The observed decrease in activity could be due to the electron deficiency of the imidazoquinoline ring system.

Some notable compounds are the C7-methoxy compounds, **4** and **5**, and the C7-hydroxyl, **14**, as these are substantially more active agonists, which suggests that the increase of electron density of the imidazoquinoline ring system may be increasing the hydrogen bonding interactions of the C4-amine and N5 pyridine nitrogen, or the increase could be contributed to a potential hydrogen bond from the neighboring tyrosine in the binding pocket.^28^

The imidazoquinolines were further screened for cytokine production in canine peripheral blood mononuclear cells (cPBMCs). The agonists at 20 μM concentrations were evaluated for TNFα, IL-12, IL-6, IL-10, IL-2, and IFNγ production by ELISA following incubation with 2 × 10^6^ PBMC/mL for 29 hours. TNFα and IL-12 were chosen as they are typically seen in a Th-1 cytokine response. None of the compounds studied activated IL-6, IL-10 or IL-2 significantly above vehicle. However, as expected nearly all compounds in this study activated IL-12 and TNFα. A trend is observed that IL-12 production increases when the tertiary alcohol of the N1 position is present, suggesting that TLR8 helps increase IL-12 production. Also having a tertiary alcohol at the N1 position appears to activate INFγ production. It is also observed that EDGs are typically stronger activators of TNFα and IL-12 when compared to EWGs. One compound that is interesting is **14**. It did not follow the trends above. It has one of the highest TNFα responses but one of the lowest IL-12 responses observed despite that it is strongly electron donating and one of the strongest activators of TLR7. Possible explanations are time courses changes in the cytokine response, or other secondary interactions of the compound within cell lines, which we will examine in future in vivo studies. Also note that we evaluated a limited panel of cytokines. While the strong IL12 response and limited activation of IL10 is strongly associated with a Th1 biased response, we do not have sufficient data to conclude none of the compounds induced a Th2 differentiation. Currently we are evaluating selected compounds for in vivo for induction of tumor-infiltration lymphocytes and natural killer cells in synergistic mouse tumors, which is more indicative of a pro-antitumor activity than the broad classification of compounds as Th1 or Th2 biased.

In this study the SAR of the C7 position of imidazoquinolines was further evaluated by synthesizing compounds **4 – 14** with overall yields around 30% at the gram scale. We observed that changes to the N1 and C2 positions combined with the C7 position maintained activity trends consistent with findings from others. We also observed that TLR7/8 tolerates all of the target C7 substitutions made in this study when the N1 and C2 are substituted appropriately. We also observed a trend that EDGs at C7 of imidazoquinolines are stronger activators of TLR7/8 when compared to EWGs. This study also found TLR7/8 activating substitutions generally increased TNFα and IL-12 cytokine responses and that TLR8 activation potentiated an IFNγ response. However, one compound, **14**, breaks this trend. Despite being one of the strongest TLR7 selective agonists **14** barely potentiates any cytokine induction. Other notable compounds produced from this study are **5**, **8**, and **14** which are up to 4-fold and 2-fold more active than resiquimod for TLR8 and/or TLR7, respectively, and up to 100x more active than the FDA approved imiquimod for TLR7. Only a few TLR7/8 agonists have been reported to date with <100nM activity, and this is the first report of high potency agonists based on modifications at the C7 position using the methoxy, chloro, and nitrile functional groups. The results in cPBMCs show that different C7 functionalities alter the agonist-cytokine response profile. Additional investigations of the C7 position are currently underway and will be disclosed in due course.

## Appendix. A Supplementary data

Supplementary data file included.

## AUTHOR INFORMATION

The authors declare no competing financial interests.

## ACKNOWLEDGMENT

The following funding sources supported this work: NIH R01CA173292 (PI:MLF); NIH0074950; NIH Biotechnology Predoctoral Training Program, NIGMS, T32-GM008359 (JRH); a gift from the Brandmeyer Family Foundation (MLF); PhRMA Foundation predoctoral fellowship (PAK). NMR instruments were acquired with support from NIH P50GM069663 and S10RR014767. We also thank Frank Forrest and Chad Groer for their gracious assistance in cytokine testing.

