## Supplemental files for "Further Exploration of the SAR of Imidazoquinolines; Identification of Potent C7 Substituted Imidazoquinolines"

**Experimental Methods**

**General Experimental Conditions.**

All solvents were ACS grade or better and used as received. All starting chemicals were purchased from AK Scientific, Ark Pharm, AstraTech, Combi-Blocks, eNovation Chemicals, Fisher, Oakwood Chemical, Sigma-Aldrich, and Strem Chemicals or were synthesized using the cited literature protocol. All reactions were conducted under an atmosphere of N_2_ or Ar unless stated otherwise. Thin layer chromatography (TLC) was performed on 0.25 mm glass-backed silica GF plates from Analtech. Developed plates were visualized with a hand-held UV lamp. Flash chromatography was performed on a CombiFlash RF system using pre-packed columns from Teledyne-Isco. ^1^H and ^13^C NMR spectra were recorded on a 500 MHz Bruker AVIII spectrometer equipped with a cryoprobe or a Bruker 400 MHz spectrometer in the noted solvent. Peaks are reported as chemical shift (δ) in ppm, coupling constants reported in (*J*) are reported in Hz, and number of protons (H) are noted from integrations in MestReNova software. High resolution mass spectra (HRMS) were obtained on an LCT Premier (Micromass Ltd., Manchester UK) time-of-flight (TOF) mass spectrometer (MS) equipped with an ESI interface. Final compounds were tested for purity and confirmed to be greater than 90% before evaluation in cell assays by HPLC/MS. Compounds **16a, 17a and 18a** were synthesized according to literature.^1^

**Preparation 7-Chloroimidazoquinolone (i)**

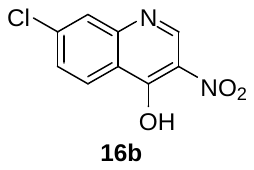

**7-Chloro-4-hydroxy-3-nitroquinoline (16b)**

The procedure was adapted from Gerster *et al.* with modifications.^1^ In a round bottom flask, 7-chloro-4-hydroxyquinoline was suspended in propionic acid and heated to reflux with a water cooled condenser open to air. 70% nitric acid (2.2 equiv.) was added dropwise over 15 minutes. The reaction was refluxed for 1 h. The reaction was allowed to cool to room temperature and diluted with EtOH and the solid was collected by vacuum filtration. The solid was washed with cold EtOH followed by hexanes, and **16b** was isolated in 70% yield as a tan solid and carried to the next step without further purification. ^1^H NMR (400 MHz, DMSO-*d*_6_) δ 13.01 (s, 1H), 8.24 (d, *J* = 8.7 Hz, 1H), 7.76 (d, *J* = 2.0 Hz, 1H), 7.55 (dd, *J* = 8.7, 2.0 Hz, 1H)).

**General Procedure A (ii & iii):**

The procedure was adapted from Gerster *et al.* with modifications.^1^ Phosphorus oxychloride (1.2 equiv.) was added dropwise to a well stirred suspension of the 7-substituted-4-hydroxy-3-nitroquinoline in anhydrous DMF (1.8 mL per mmol) and an exothermic reaction was observed. After addition of POCl_3_ was completed, the reaction was heated at 50 °C and stirred for 30 min. The resulting solution was cooled to room temperature and poured into ice water (7.5 mL per mmol). The resulting solid was collected by vacuum filtration, washed with water and pressed dry. The moist solid was added to a round bottom flask and suspended in EtOH (5 mL per mmol), Et_3_N (2 equiv.), and the appropriate alkyl amine (1.3 equiv.) and refluxed for 15 min. Water was added to the solution and the solid was collected by vacuum filtration and carried onto the next step without further purification.

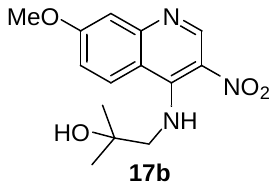

**4-(2-Hydroxy-2-methylproylamino)-7-methoxy-3-nitroquinoline (17b)**

The title compound was prepared according to the general procedure A using **16a** and 1-Amino-2-methyl-2-propanol to obtain a bright yellow solid in 80% yield. ^1^H NMR (400 MHz, Methanol-*d*_4_) δ 9.20 (s, 1H), 8.38 (d, *J =* 9.5 Hz, 1H), 7.28 (d, *J =* 2.7 Hz, 1H), 7.19 (dd, *J =* 9.4, 2.7 Hz, 1H), 4.59 (s, 1H), 3.97 (s, 3H), 3.91 (s, 2H), 1.29 (s, 6H).

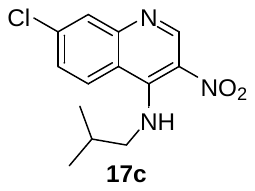

**7-Chloro-4-(2-methylpropylamino)-3-nitroquinoline (17c)**

The title compound was prepared according to the general procedure A using **16b** and isobutylamine to obtain a bright yellow solid in 79% yield. ^1^H NMR (CDCl_3_, 400 MHz) δ 9.87 (s, 1H), 9.37 (s, 1H), 8.25 (d, *J* = 9.0 Hz, 1H), 8.03 (d, *J* = 2.2 Hz, 1H), 7.45 (dd, *J* = 9.1, 2.2 Hz, 1H), 3.77 (dd, *J* = 6.5, 4.8 Hz, 2H), 2.09 (dt, *J* = 13.3, 6.6 Hz, 1H), 1.11 (d, *J* = 6.7 Hz, 6H).

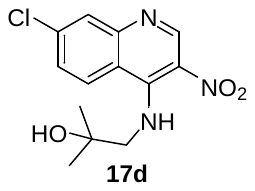

**7-Chloro-4-(2-hydroxy-2-methylpropylamino)-3-nitroquinoline (17d)**

The title compound was prepared according to the general procedure A using **16b** and 1-Amino-2-methyl-2-propanol to obtain a bright yellow solid in 77% yield. ^1^H NMR (CDCl_3_, 400 MHz) δ 9.93 (s, 1H), 9.37 (s, 1H), 8.18 (d, *J =* 9.0 Hz, 1H), 7.99 (d, *J =* 2.4 Hz, 1H), 7.44 (dd, *J =* 9.1, 2.3 Hz, 1H), 3.85 (d, *J =* 4.9 Hz, 2H), 1.36 (s, 6H).

#### **General Procedure B (iv & v):**

A suspension of 7-substituted amino nitroquinoline in EtOH was heated to reflux in open air. Iron (dust 5 equiv.) was added followed by aqueous NH_4_Cl (2.3 M, 5 equiv.) and refluxed for 2 hours. The solution was allowed to cool and was filtered through a Celite plug and eluted with EtOAc. The volume of solvent was reduced on a rotovap and basified with saturated Na_2_CO_3_. The aqueous layer was extracted with EtOAc (3x). The combined organic layers were dried with Na_2_SO_4_, filtered, and evaporated on a rotovap to dryness. Valeric or ethoxy acetic acid (10 equiv.) was added to diaminoquinolines in a small round bottom flask. The suspension was heated to reflux (~150 °C) in open air until water ceased to be released from the reaction. The reaction was allowed to cool and diluted with water then basified with 6M NaOH. The water was extracted with DCM (3x) and the combined organic layer was washed with brine and dried with Na_2_SO_4_, filtered, and concentrated on rotovap. The compound was purified by CombiFlash (0-10% MeOH in DCM).

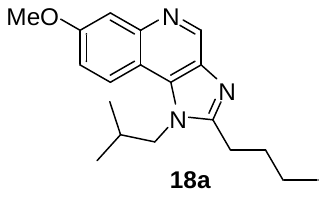

**2-Butyl-7-methoxy-1-(2-methylpropyl)-1*H*-imidazo[4,5-c]quinoline (18a)**

The title compound was prepared according to the general procedure C using **17a** to obtain a brown solid in 95% yield. ^1^H NMR (400 MHz, CDCl_3_) δ 9.21 (s, 1H), 7.98 (d, *J =* 9.2 Hz, 1H), 7.65 (d, *J =* 2.7 Hz, 1H), 7.29 – 7.26 (m, 1H), 4.27 (d, *J =* 7.6 Hz, 2H), 3.96 (s, 3H), 3.03– 2.88 (m, 2H), 2.40 – 2.27 (m, 1H), 1.94 (m, 2H), 1.50 (m, 2H), 1.01 (m, 9H).

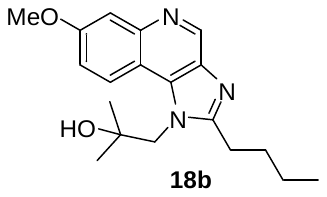

**2-Butyl-1-(2-hydroxy-2-methylpropyl)-7-methoxy-1*H*-imidazo[4,5-c]quinoline (18b)**

The title compound was prepared according to the general procedure C using **17b** to obtain a brown solid in 48% yield. ^1^H NMR (400 MHz, CDCl_3_) δ 8.94 (s, 1H), 8.53 (s, 1H), 7.81 (s, 1H), 7.33 (d, *J =* 9.4 Hz, 1H), 4.62 (s, 2H), 3.97 (s, 1H), 3.88 (s, 3H), 3.06 (t, 2H), 1.95 (p, *J =* 7.8 Hz, 2H), 1.52 (q, *J =* 7.5 Hz, 2H), 1.47 (s, 6H), 1.01 (t, *J =* 7.3 Hz, 2H).

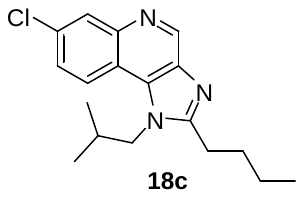

**2-Butyl-7-chloro-1-(2-methylpropyl)-1*H*-imidazo[4,5-c]quinoline (18c)**

The title compound was prepared according to the general procedure C using **17c** to obtain a brown solid in 80% yield. ^1^H NMR (CDCl_3_, 500 MHz) δ 9.27 (s, 1H), 8.38 (d, *J =* 2.2 Hz, 1H), 8.05 (d, *J =* 9.0 Hz, 1H), 7.62 (dd, *J =* 9.0, 2.2 Hz, 1H), 4.32 (d, *J =* 7.5 Hz, 2H), 2.99 – 2.93 (m, 2H), 2.33 (m, 1H), 1.96 (m, 2H), 1.52 (m, 2H), 1.03 (d, *J =* 6.7 Hz, 6H), 1.02 (t, *J =* 7.4 Hz, 3H).

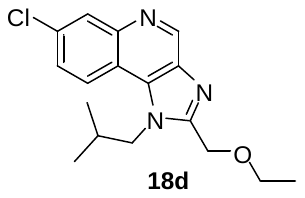

**7-Chloro-2-(ethoxymethyl)-1-(2-methylpropyl)-1*H*-imidazo[4,5-c]quinoline (18d)**

The title compound was prepared according to the general procedure C using **17c** to obtain a brown solid in 88% yield. ^1^H NMR (400 MHz, CDCl_3_) δ 9.30 (s, 1H), 8.35 (d, *J =* 2.z2 Hz, 1H), 8.07 (d, *J =* 9.0 Hz, 1H), 7.64 (dd, *J =* 8.9, 2.2 Hz, 1H), 4.89 (s, 2H), 4.51 (d, *J =* 7.7 Hz, 2H), 3.61 (q, *J =* 7.0 Hz, 2H), 2.37 (m, 1H), 1.25 (t, *J =* 7.0 Hz, 3H), 1.09 – 1.00 (d, *J* = 6.7 Hz, 6H).

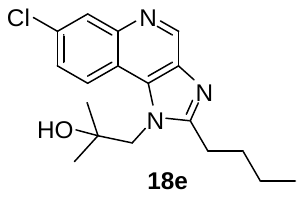

**2-Butyl-7-chloro-1-(2-hydroxy-2-methylpropyl)-1*H*-imidazo[4,5-c]quinoline (18e)**

The title compound was prepared according to the general procedure C using **17d** to obtain a brown solid in 87% yield. ^1^H NMR (400 MHz, CDCl_3_) δ 8.99 (s, 1H), 8.52 (d, *J =* 9.0 Hz, 1H), 8.13 (s, 1H), 7.55 (dd, *J =* 9.1, 2.2 Hz, 1H), 4.62 (s, 2H), 3.06 (s, 2H), 1.95 (p, *J =* 7.7 Hz, 2H), 1.51 (m 2H), 1.46 (s, 6H), 1.01 (t, *J =* 7.3 Hz, 3H).

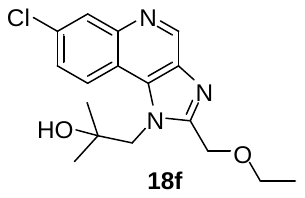

**7-Chloro-2-(ethoxymethyl)-1-(2-hydroxy-2-methylpropyl)-1*H*-imidazo[4,5-c]quinoline (18f)**

The title compound was prepared according to the general procedure C using **17d** to obtain a brown solid in 87% yield. ^1^H NMR (400 MHz, CDCl_3_) δ 9.14 (s, 1H), 8.41 (s, 1H), 8.25 (s, 1H), 7.60 (d, *J =* 8.9 Hz, 1H), 4.99 (s, 2H), 4.83 (s, 2H), 3.65 (q, *J =* 7.0 Hz, 2H), 1.40 (s, 6H), 1.25 (dd, *J =* 7.4, 6.7 Hz, 3H).

#### **General Procedure C (vi):**

The procedure was adapted from Gerster *et al.* with modifications.^1^ In a round bottom flask, imidazoquinoline was dissolved in CHCl_3_ and stirred at room temperature. *m*CPBA (1 + 1 equiv.) was added portion-wise 1 h apart. After 3 hours the reaction was diluted with DCM, washed with saturated Na_2_CO_3_ aqueous solution, extracted with DCM (3x), dried with Na_2_SO_4_, filtered, and concentrated on rotovap. The compound was purified by CombiFlash (0-10% MeOH in DCM).

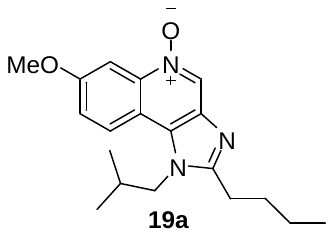

**2-Butyl-7-methoxy-1-(2-methylpropyl)-1*H*-imidazo[4,5-c]quinoline 5-oxide (19a)**

The title compound was prepared according to the general procedure D using **18a** to obtain a brown solid in 45% yield. ^1^H NMR (DMSO-*d*_6_, 500 MHz) δ 8.99 (s, 1H), 8.24 (d, *J =* 9.0 Hz, 1H), 8.22 (d, *J =* 2.8 Hz, 1H), 7.47 (dd, *J =* 9.2, 2.9 Hz, 1H), 4.37 (d, *J =* 7.6 Hz, 2H), 2.91 (t, *J =* 7.6 Hz, 2H), 2.13 (dt, *J =* 13.5, 6.8 Hz, 1H), 1.82 (p, *J =* 7.5 Hz, 2H), 1.44 (h, *J =* 7.4 Hz, 2H), 0.94 (t, *J =* 7.5 Hz, 3H), 0.91 (d, *J =* 6.7 Hz, 6H). HRMS (*m/z*): [M+H]^+^ calcd for C_19_H_26_N_3_O_2_, 328.2025; found 328.2034.

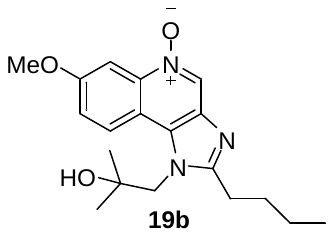

**2-Butyl-1-(2-hydroxy-2-methylpropyl)-7-methoxy-1*H*-imidazo[4,5-c]quinoline 5-oxide (19b)**

The title compound was prepared according to the general procedure D using **18b** to obtain a brown solid in 90% yield. ^1^H NMR (400 MHz, CDCl_3_) δ 8.67 (d, *J =* 9.3 Hz, 1H), 8.02 (s, 1H), 7.52 (d, *J =* 2.7 Hz, 1H), 7.19 (dd, *J =* 9.3, 2.8 Hz, 1H), 4.10 (s, 1H), 3.83 (s, 3H), 3.43 (d, *J =* 3.8 Hz, 1H), 2.92 (s, 3H), 1.86 (q, *J =* 7.7 Hz, 2H), 1.30 (d, *J =* 2.0 Hz, 2H), 0.98 (t, *J =* 7.4 Hz, 3H). HRMS (*m/z*): [M+H]^+^ calcd for C_19_H_26_N_3_O_3_, 344.1974; found 344.1986.

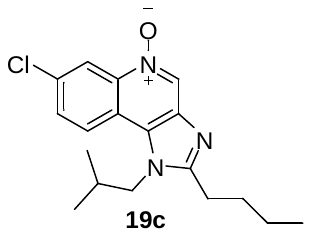

**2-Butyl-7-chloro-1-(2-methylpropyl)-1*H*-imidazo[4,5-c]quinoline 5-oxide (19c)**

The title compound was prepared according to the general procedure D using **18c** to obtain a brown solid in 80% yield. ^1^H NMR (CDCl_3_, 500 MHz) δ 9.58 (s, 1H), 8.88 (d, *J =* 2.0 Hz, 1H), 8.20 (d, *J =* 9.0 Hz, 1H), 7.86 (dd, *J =* 9.0, 2.2 Hz, 1H), 4.39 (d, *J =* 7.6 Hz, 2H), 3.04 – 2.97 (m, 2H), 2.30 (dt, *J =* 13.8, 6.9 Hz, 1H), 1.97 (p, *J =* 7.6 Hz, 2H), 1.52 (ddd, *J =* 14.2, 7.2, 5.4 Hz, 3H), 1.07 (d, *J =* 6.7 Hz, 6H), 1.03 (t, *J =* 7.3 Hz, 3H); HRMS (*m/z*): [M+Na]^+^ calcd for C_18_H_22_ClN_3_ONa, 354.1349; found 354.1362.

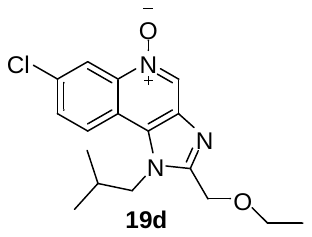

**7-Chloro-2-(ethoxymethyl)-1-(2-methylpropyl)-1*H*-imidazo[4,5-c]quinoline 5-oxide (19d)**

The title compound was prepared according to the general procedure D using **18d** to obtain a brown solid in 76% yield. ^1^H NMR (400 MHz, DMSO-*d*_6_) δ 9.12 (d, *J =* 9.9 Hz, 1H), 8.83 – 8.73 (m, 1H), 8.40 (dd, *J =* 20.5, 9.0 Hz, 1H), 7.92 (ddd, *J =* 10.9, 8.9, 2.4 Hz, 1H), 4.80 (s, 2H), 4.55 – 4.43 (m, 2H), 3.59 (q, *J =* 7.0 Hz, 2H), 2.22 (p, *J =* 7.1 Hz, 1H), 1.16 (t, *J =* 7.0 Hz, 3H), 0.92 (d, *J =* 6.7 Hz, 6H). HRMS (*m/z*): [M+H]^+^ calcd for C_17_H_21_ClN_3_O_2_, 334.1322; found 334.1328.

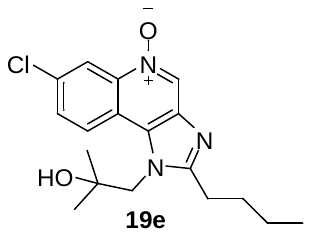

**2-Butyl-7-chloro-1-(2-hydroxy-2-methylpropyl)-1*H*-imidazo[4,5-c]quinoline 5-oxide (19e)**

The title compound was prepared according to the general procedure D using **18e** to obtain a brown solid in 71% yield. ^1^H NMR (400 MHz, CDCl_3_) δ 9.04 (s, 1H), 8.72 (d, *J =* 9.1 Hz, 1H), 8.48 (s, 1H), 7.65 (d, *J =* 8.5 Hz, 1H), 4.48 (s, 2H), 3.01 (s, 2H), 1.93 (p, *J =* 7.7 Hz, 2H), 1.51 (dt, *J =* 15.0, 7.4 Hz, 2H), 1.44 (s, 6H), 1.23 (d, *J =* 16.9 Hz, 1H), 1.01 (t, *J =* 7.3 Hz, 3H). HRMS (*m/z*): [M+H]^+^ calcd for C_18_H_23_ClN_3_O_2_, 348.1479; found 348.1468.

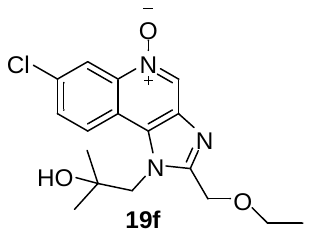

**7-Chloro-2-(ethoxymethyl)-1-(2-hydroxy-2-methylpropyl)-1*H*-imidazo[4,5-c]quinoline**

**5-oxide (19f)**

The title compound was prepared according to the general procedure D using **18f** to obtain a brown solid in 99% yield. ^1^H NMR (CDCl_3_, 500 MHz) δ 9.18 (s, 1H), 8.66 (d, *J =* 8.9 Hz, 1H), 8.64 – 8.60 (m, 1H), 7.73 (d, *J =* 8.3 Hz, 1H), 4.97 (s, 2H), 4.80 (s, 2H), 3.61 (q, *J =* 7.0 Hz, 2H), 1.41 (s, 6H), 1.24 (t, *J =* 7.0 Hz, 3H). HRMS (*m/z*): [M+Na]^+^ calcd for C_17_H_20_ClN_3_O_3_Na, 372.1091; found 372.1085.

#### **General Procedure D (vii)**:

The procedure was adapted from Gerster *et al.* with modifications.^1^ Concentrated NH_4_OH (1 mL per mmol) was added to a round bottom flask containing N-oxide imidazoquinoline (1 equiv.) in anhydrous DCM (1 mL per mmol) stirring vigorously at room temperature. *p*-TsCl (1.1 equiv.) was added to an addition funnel and dissolved in anhydrous DCM (1 mL per mmol). The *p*-TsCl in DCM was added dropwise slowly over 15 min minutes and more DCM was added to help push residual *p*-TsCl into the flask to the round bottom. An exotherm was observed during the addition. The reaction was allowed to stir for another 15 min after addition of *p*-TsCl was completed. Water was added and if precipitate formed, the solid was isolated by vacuum filtration. The mother liquor or biphasic mixture was extracted with DCM, washed with brine, dried with Na_2_SO_4_, and concentrated *in vacuo* and purified by CombiFlash.

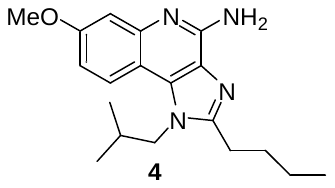

**4-Amino-2-butyl-7-methoxy-1-(2-methylpropyl)-1*H*-imidazo[4,5-c]quinoline (4)**

The title compound was prepared according to the general procedure E using **19a** to obtain a brown solid in 77% yield. ^1^H NMR (CDCl_3_, 500 MHz) δ 7.76 (d, *J =* 9.0 Hz, 1H), 7.30 (d, *J =* 2.7 Hz, 1H), 7.00 (dd, *J =* 9.0, 2.6 Hz, 1H), 6.43 (s, 2H), 4.20 (d, *J =* 7.6 Hz, 2H), 3.92 (s, 3H), 2.90 – 2.83 (m, 2H), 2.31 (dp, *J =* 13.4, 6.6 Hz, 1H), 1.86 (p, *J =* 7.6 Hz, 2H), 1.49 (h, *J =* 7.4 Hz, 2H), 1.02 – 0.97 (m, 9H); ^13^C NMR (CDCl_3_, 126 MHz) δ 159.49, 154.41, 150.80, 134.77, 124.73, 121.28, 114.23, 108.69, 105.40, 55.70, 52.71, 30.01, 29.28, 27.58, 22.73, 19.92, 19.79, 13.99. HRMS (*m/z*): [M+H]^+^ calcd for C_19_H_27_N_4_O, 327.2185; found 327.2196.

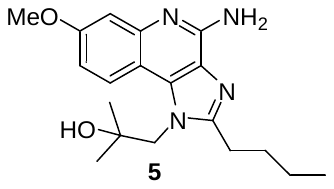

**4-Amino-2-butyl-1-(2-hydroxy-2-methylpropyl)-7-methoxy-1*H*-imidazo[4,5-c]quinoline (5)**

The title compound was prepared according to the general procedure E using **19b** to obtain a brown solid in 71% yield. ^1^H NMR (500 MHz, DMSO-*d*_6_) δ 9.12 (d, *J =* 10.7 Hz, 1H), 8.80 (dd, *J =* 5.0, 2.3 Hz, 1H), 8.40 (dd, *J =* 20.5, 9.1 Hz, 1H), 7.91 (ddd, *J =* 11.1, 9.0, 2.4 Hz, 1H), 4.80 (s, 1H), 4.49 (s, *J =* 21.9, 7.7 Hz, 2H), 3.59 (q, *J =* 7.0 Hz, 2H), 2.22 (p, *J =* 7.0 Hz, 1H), 1.16 (t, *J =* 7.0 Hz, 3H), 0.92 (d, *J =* 6.6 Hz, 6H); ^13^C NMR (126 MHz, DMSO-*d*_6_) δ 157.65, 154.34, 151.79, 146.23, 134.09, 124.72, 122.33, 110.62, 109.58, 106.88, 70.79, 54.87, 54.46, 48.59, 29.80, 26.76, 22.05, 13.87. HRMS (*m/z*): [M+H]^+^ calcd for C_19_H_27_N_4_O_2_, 343.2134; found 343.2129.

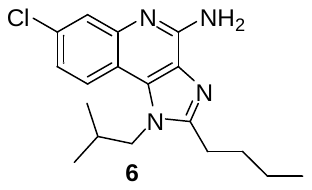

**4-Amino-2-butyl-7-chloro-1-(2-methylpropyl)-1*H*-imidazo[4,5-c]quinoline (6)**

The title compound was prepared according to the general procedure E using **19c** to obtain a brown solid in 81% yield. ^1^H NMR (DMSO-*d*_6_, 500 MHz) δ 7.97 (d, *J =* 8.9 Hz, 1H), 7.56 (d, *J =* 2.3 Hz, 1H), 7.26 (dd, *J =* 8.8, 2.3 Hz, 1H), 6.69 (s, 2H), 4.33 (d, *J =* 7.6 Hz, 2H), 2.93 – 2.87 (m, 2H), 2.11 (dq, *J =* 13.8, 6.9 Hz, 1H), 1.85 – 1.76 (m, 2H), 1.44 (dt, *J =* 14.7, 7.4 Hz, 2H), 0.95 (t, *J =* 7.4 Hz, 3H), 0.91 (d, *J =* 6.6 Hz, 6H); ^13^C NMR (DMSO-*d*_6_, 126 MHz) δ 153.99, 152.59, 145.83, 132.08, 130.51, 126.58, 124.81, 121.97, 120.90, 113.57, 51.31, 29.65, 28.82, 26.43, 21.93, 19.16, 13.81. HRMS (*m/z*): [M+H]^+^ calcd for C_18_H_24_ClN_4_, 331.1689; found 331.1702.

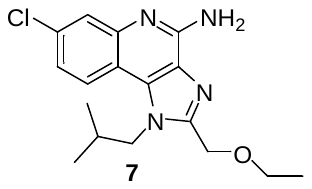

**4-Amino-7-chloro-2-(ethoxymethyl)-1-(2-methylpropyl)-1*H*-imidazo[4,5-c]quinoline (7)**

The title compound was prepared according to the general procedure E using **19d** to obtain a brown solid in 78% yield. ^1^H NMR (500 MHz, DMSO-*d*_6_) δ 8.01 (d, *J =* 8.8 Hz, 1H), 7.58 (d, *J =* 2.3 Hz, 1H), 7.27 (dd, *J =* 8.8, 2.3 Hz, 1H), 6.88 (s, 2H), 4.75 (s, 2H), 4.42 (d, *J =* 7.7 Hz, 2H), 3.56 (q, *J =* 7.0 Hz, 2H), 2.20 (hept, *J =* 6.9 Hz, 1H), 1.15 (t, *J =* 7.0 Hz, 3H), 0.91 (d, *J =* 6.6 Hz, 6H); ^13^C NMR (126 MHz, DMSO-*d*_6_) δ 152.93, 149.72, 146.29, 132.82, 131.12, 126.48, 124.82, 122.43, 121.00, 113.45, 65.41, 64.16, 51.77, 28.61, 19.25, 14.92. HRMS (*m/z*): [M+H]^+^ calcd for C_17_H_22_ClN_4_O, 333.1482; found 333.1475.

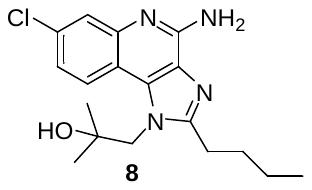

**4-Amino-2-butyl-7-chloro-1-(2-hydroxy-2-methylpropyl)-1*H*-imidazo[4,5-c]quinoline (8)**

The title compound was prepared according to the general procedure E using **19e** to obtain a brown solid in 73% yield. ^1^H NMR (CDCl_3_, 500 MHz) δ 8.29 (d, *J =* 8.9 Hz, 1H), 7.53 (d, *J =* 2.3 Hz, 1H), 7.17 (dd, *J =* 8.8, 2.3 Hz, 1H), 6.63 (s, 2H), 4.77 (s, 1H), 4.51 (s, 2H), 3.00 (t, 2H), 1.78 (ddd, *J =* 15.4, 8.9, 7.1 Hz, 2H), 1.42 (h, *J =* 7.4 Hz, 2H), 1.16 (s, 6H), 0.94 (t, *J =* 7.4 Hz, 3H); ^13^C NMR (CDCl_3_, 126z MHz) δ 155.39, 152.57, 145.93, 133.30, 130.16, 126.28, 124.58, 123.17, 120.10, 114.26, 70.76, 54.49, 48.59, 29.78, 26.80, 22.03, 13.86. HRMS (*m/z*): [M+H]^+^ calcd for C_18_H_24_ClN_4_O, 347.1639; found 347.1649.

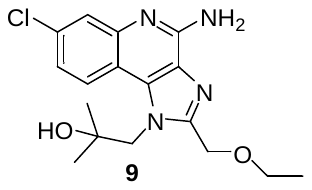

**4-Amino-7-chloro-2-(ethoxymethyl)-1-(2-hydroxy-2-methylpropyl)-1*H*-imidazo[4,5-c]quinoline (9)**

The title compound was prepared according to the general procedure E using **19f** to obtain a brown solid in 84% yield. ^1^H NMR (DMSO-*d*_6_, 500 MHz) δ 8.30 (d, *J =* 8.9 Hz, 1H), 7.54 (d, *J =* 2.3 Hz, 1H), 7.19 (dd, *J =* 8.8, 2.3 Hz, 1H), 6.79 (s, 2H), 4.88 (s, 1H), 4.64 (s, 2H), 3.51 (q, *J =* 7.0 Hz, 2H), 1.16 (s, 6H), 1.13 (t, *J =* 7.0 Hz, 3H); ^13^C NMR (DMSO-*d*_6_, 126 MHz) δ 153.38, 151.61, 146.91, 134.40, 131.26, 126.67, 125.14, 123.86, 120.75, 114.52, 71.11, 65.80, 65.28, 55.25, 15.47. HRMS (*m/z*): [M+H]^+^ calcd for C_17_H_22_ClN_4_O_2_, 349.1431; found 349.1425.

**Preparation of Aryl Nitriles (xii):**

**General Procedure F**

The procedure was adapted from Littke *et al.* with modifications.^2^ Aryl chloride (1 equiv.), Pd(TFA)_2_ (5%), Zn (dust, 20%), rac-2-(Di-tert-butylphosphino)-1,1′-binaphthyl (TrixiePhos) (10%), Zn(CN)_2_ (56%) were added to an oven-dried 40 mL vial and vacuum flushed with nitrogen three times. Anhydrous dimethylacetamide (DMAC) (0.19 M) was added via syringe to the sealed vial. The vial was shaken and placed on a heating mantel at 95 °C and stirred overnight. The mixture was allowed to cool to room temperature and filtered through a plug of Celite using either 100% EtOAc or 20% MeOH in DCM to elute product. The solvent was then concentrated down and redissovled in EtOAc and washed with water (4-5x by 10x the volume of DMAC), washed with brine and dried with Na_2_SO_4_. Solvent was concentrated down on rotovap and purified by CombiFlash (0-10% MeOH in DCM).

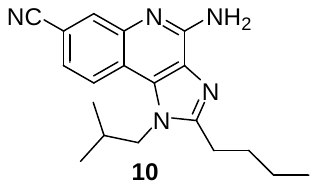

**4-Amino-2-butyl-7-carbonitrile-1-(2-methylpropyl)-1*H*-imidazo[4,5-c]quinoline (10)**

The title compound was prepared according to the general procedure F using **6** to obtain a brown solid in 82% yield. ^1^H NMR (500 MHz, DMSO-*d*_6_) δ 8.13 (d, *J* = 8.5 Hz, 1H), 7.98 (t, *J* = 1.5 Hz, 1H), 7.56 (dt, *J* = 8.3, 1.5 Hz, 1H), 6.90 (s, 2H), 4.38 (d, *J* = 7.6 Hz, 2H), 2.94 (t, *J* = 6.3 Hz, 2H), 2.18 – 2.06 (m, 1H), 1.83 (t, *J* = 7.7 Hz, 2H), 1.46 (q, *J* = 7.6 Hz, 2H), 0.98 – 0.94 (t, 3H), 0.93 (d, *J* = 6.6 Hz, 7H); ^13^C NMR (500 MHz, DMSO-*d*_6_) δ 155.29, 152.90, 144.07, 131.51, 130.47, 128.12, 122.36, 121.70, 119.40, 118.00, 108.27, 51.38, 29.60, 28.90, 26.48, 21.91, 19.13, 13.80. HRMS (*m/z*): [M+H]^+^ calcd for C_19_H_24_N_5_, 322.2032; found 322.2034.

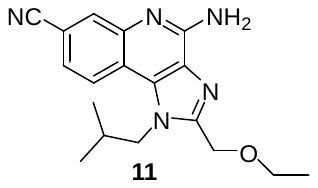

**4-Amino-7-carbonitrile-2-(ethoxymethyl)-1-(2-methylpropyl)-1*H*-imidazo[4,5-c]quinoline (11)**

The title compound was prepared according to the general procedure F using **7** to obtain a brown solid in 80% yield. ^1^H NMR (400 MHz, CDCl_3_) δ 8.22 (d, *J* = 1.7 Hz, 1H), 8.01 (d, *J* = 8.5 Hz, 1H), 7.61 (dd, *J* = 8.5, 1.7 Hz, 1H), 6.42 (s, 2H), 4.86 (s, 2H), 4.48 (d, *J* = 7.7 Hz, 2H), 4.15 (q, *J* = 7.1 Hz, 1H), 3.65 (q, *J* = 7.0 Hz, 2H), 2.36 (dt, *J* = 13.9, 7.0 Hz, 1H), 1.31 – 1.24 (m, 3H), 1.07 (d, *J* = 6.7 Hz, 6H); ^13^C NMR (126 MHz, DMSO-*d*_6_) δ 151.85, 149.18, 143.94, 132.94, 132.40, 129.13, 124.57, 122.81, 119.48, 119.06, 109.77, 66.15, 64.69, 52.50, 29.12, 19.74, 15.45. HRMS (*m/z*): [M+H]^+^ calcd for C_18_H_22_N_5_O, 324.1824; found 324.1816.

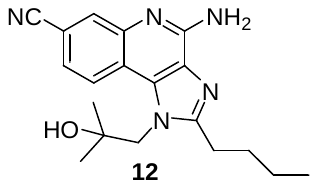

**4-Amino-2-butyl-7-carbonitrile-1-(2-hydroxy-2-methylpropyl)-1*H*-imidazo[4,5-c]quinoline (12)**

The title compound was prepared according to the general procedure F using **8** to obtain a brown solid in 81% yield. ^1^H NMR (500 MHz, DMSO-*d*_6_) δ 8.45 (d, *J =* 8.6 Hz, 1H), 7.94 (d, *J =* 1.8 Hz, 1H), 7.46 (dd, *J =* 8.6, 1.8 Hz, 1H), 6.85 (s, 2H), 4.78 (s, 1H), 4.52 (s, 2H), 3.02 (t, *J =* 7.8 Hz, 2H), 1.80 (ddd, *J =* 15.3, 8.9, 7.1 Hz, 2H), 1z.42 (h, *J =* 7.4 Hz, 2H), 1.17 (s, 6H), 0.94 (t, *J =* 7.4 Hz, 3H); ^13^C NMR (126 MHz, DMSO-*d*_6_) δ 156.64, 152.92, 144.16, 132.82, 130.29, 127.82, 123.04, 121.50, 119.52, 118.79, 107.94, 70.75, 54.55, 29.75, 26.88, 22.02, 13.86. HRMS (*m/z*): [M+H]^+^ calcd for C_19_H_24_N_5_O, 338.1981; found 338.1981.

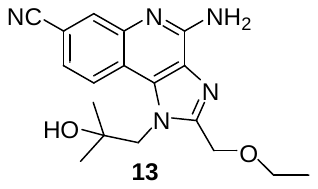

**4-Amino-7-carbonitrile-2-(ethoxymethyl)-1-(2-hydroxy-2-methylpropyl)-1*H*-imidazo[4,5-c]quinoline (13)**

The title compound was prepared according to the general procedure F using **9** to obtain a brown solid in 76% yield. ^1^H NMR (500 MHz, DMSO-*d*_6_) δ 8.47 (d, *J =* 8.6 Hz, 1H), 7.96 (d, *J =* 1.8 Hz, 1H), 7.48 (dd, *J =* 8.5, 1.8 Hz, 1H), 7.01 (s, 2H), 4.89 (s, 1H), 4.68 (s, 2H), 3.52 (q, *J =* 7.0 Hz, 2H), 1.19 – 1.15 (m, 2H), 1.13 (t, *J =* 7.0 Hz, 3H); ^13^C NMR (126 MHz, DMSO-*d*_6_) δ 153.22, 152.28, 144.67, 133.41, 130.35, 127.66, 123.24, 121.66, 119.38, 118.59, 108.61, 70.61, 65.43, 64.75, 54.86, 27.58, 14.98. HRMS (*m/z*): [M+H]^+^ calcd for C_18_H_22_N_5_O_2_, 340.1774; found 340.1778.

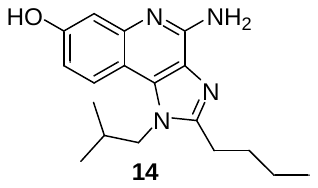

**4-Amino-2-butyl-7-hydroxy-1-(2-methylpropyl)-1*H*-imidazo[4,5-c]quinoline (14)**

**4** (1 equiv.) was added to a melt of pyridine hydrochloride (10 equiv.) and heated at 210 °C for 30 minutes. The reaction was allowed to cool for a few minutes and diluted with water then acidified with 6 N HCl to pH 1. The solid was collected by vacuum filtration and washed with hexanes and dried in air to obtain **4** in 85% yield as a light brown solid. ^1^H NMR (500 MHz, DMSO-*d*_6_) δ 14.00 (s, 1H), 10.69 (s, 1H), 8.60 (s, 3H), 7.98 (d, *J =* 9.1 Hz, 1H), 7.19 (d, *J =* 2.4 Hz, 1H), 7.09 (dd, *J =* 9.0, 2.4 Hz, 1H), 5.10 (s, 4H), 4.35 (d, *J =* 7.6 Hz, 2H), 2.92 (t, *J =* 7.6 Hz, 2H), 2.09 (dq, *J =* 13.8, 6.9 Hz, 1H), 1.81 (p, *J =* 7.6 Hz, 2H), 1.44 (h, *J =* 7.4 Hz, 2H), 0.97 – 0.90 (m, 9H); ^13^C NMR (126 MHz, DMSO-*d*_6_) δ 158.73, 155.98, 148.63, 135.85, 135.55, 123.38, 122.04, 115.11, 105.18, 102.76, 51.49, 29.25, 28.73, 26.38, 21.84, 19.11, 13.81. HRMS (*m/z*): [M+H]^+^ calcd for C_18_H_25_N_4_O 313.2028; found 313.2041.

**SEAP Reporter Assay for TLR-7/8 Activity:**

TLR7/8 analog activities were assessed using stably transfected HEK-Blue hTLR7, hTLR8, Null1-k and Null1 cell lines (InvivoGen). HEK-Blue hTLR7/8 cells are HEK293 cells co-transfected with the hTLR7/8 gene and an optimized SEAP reporter gene. HEK-Blue Null1-k and Null1 cells are transfected with the SEAP reporter gene but do not express the hTLR7/8 gene and are used as a control. None of the compounds tested showed activity in the Null1-k and Null1 controls (see KU Scholarworks), indicating no substantial NF-κB induction from endogenous, low level TLR expression by HEK293 cells.

The experiments followed manufacturer protocol as follows: SEAP levels were measured via absorbance at 637 nm using HEK-Blue Detection cell culture medium (InvivoGen), which contains a SEAP-specific colorimetric substrate. Compound stock solutions were first diluted in sterile DMSO at a concentration of 10 mM. Lower concentrations were prepared by serial dilution of the initial 10 mM stock samples into sterile DMSO and these solutions (22 μL) were subsequently diluted into sterile H_2_O (198 µL) using a BioTek Precision XS pipetting system. HEK-Blue hTLR7/8, Null1-k, and Null1 cell suspensions in sterile HEK-Blue Detection medium were prepared at a density of ~220,000 cells per mL. Clear bottom, 96-well tissue culture treated plates (Corning) were seeded with hTLR7 and Null1-k or hTLR8 and Null1 cell suspensions (180 μL/well), after which 20 μL of the various sample concentrations in 10% v/v DMSO/H_2_O were added (20 μL) in quadruplicate wells to yield the desired testing concentration in 1% v/v DMSO cell suspension. Plates were incubated at 37 °C and 5% CO_2_ in the dark. Absorbance measurements were taken at 637 nm at 12 hours, 4 replicates per concentration point, per cell line (hTLR7, hTLR8, Null1-k, Null1). Three independent measurements on different days were performed for each analog. Positive and negative controls were resiquimod and DMSO, respectively, and were tested alongside analogs 4-14 during independent measurements.

Corresponding well absorbance values for each concentration tested were averaged with respect to days tested, and the average relative NF-κB induction (Abs at 637 nm) of the null cell lines (Null1-k or Null1) were subtracted from the respective average TLR expressing cell line (hTLR7 or hTLR8) absorbance averages. The absorbance data observed was then normalized to that day’s respective Resiquimod controls based on 100% activity to better compare the biological replicates. The three independent, normalized averages for each analog were then pooled, and the data was fit using the Hill-Slope model (GraphPad Prism v7.0) using the non-linear fit from the [Agonist] vs. response – Variable Slope (four parameters) with a hill slope constrained to less than 3. These gave the dose response curves for TLR7 (**SI Figure 1**) and TLR 8 (**SI Figure 2**), along with the EC_50_ values reported in **Table 1** of the manuscript.

**Cytokine Secretion from Canine PBMCs Following TLR Agonist Stimulation:**

Peripheral blood from a healthy canine donor was obtained with appropriate informed consent of the owner and under approved institutional guidelines. Blood (8 mL per tube) was drawn via sterile venipuncture into a BD Vacutainer cell preparation tube containing sodium heparin (BD Biosciences). PBMCs were isolated within one hour following procurement by density gradient centrifugation according to the manufacturer protocol and resuspended to 2.22x10^6^ cells/mL in sterile RPMI 1640 medium with L-glutamine (Gibco) supplemented with 100 U/mL penicillin, 100 µg/mL streptomycin (Corning) and 10% (v/v) heat-inactivated Fetalgro (RMBIO). Corning 3917, 96-well tissue culture treated plates were seeded with the PBMC suspension (180 μL/well), 20 μL of the serially diluted TLR agonists were added in triplicate to yield the desired concentrations in a 1% (v/v) DMSO cell suspension. Cells were incubated at 37 °C in 5% CO_2_ for 9 hours (**SI Figure 3**) or 29 hours (**Table 1**) after which supernatants were removed and frozen at -80°C. Cytokine secretion levels were measured by ELISA (Duoset, R&D Systems) according to the manufacturer protocol. Samples and standards (n=3) were thawed and plated onto a Maxisorp flat-bottom, 384-well plate (Thermo Scientific) using a BioTek Precision XS pipetting system. Cytokine concentrations were determined by a hyperbolic or four-parameter logistic non-linear regression of the recombinant standards and subsequent interpolation using Graphpad Prism 7.0.

**SI Figure 1. Dose Response Curve of TLR7 Data**

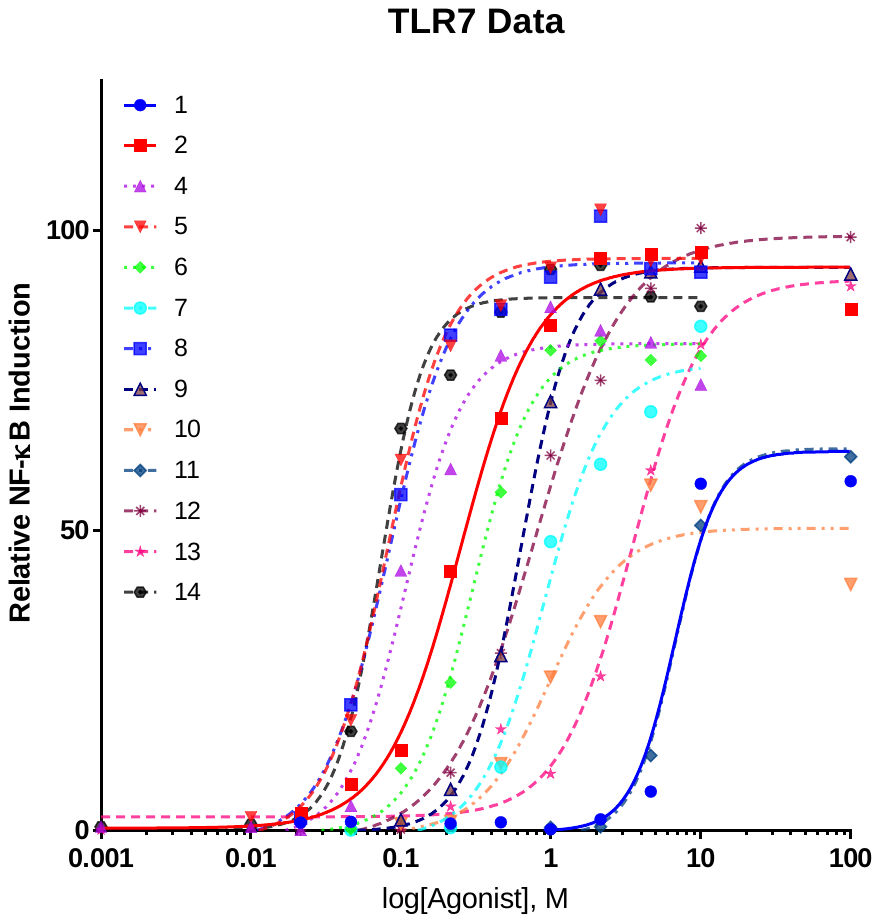

**SI Figure 2. Dose Response Curve for TLR8 Data**

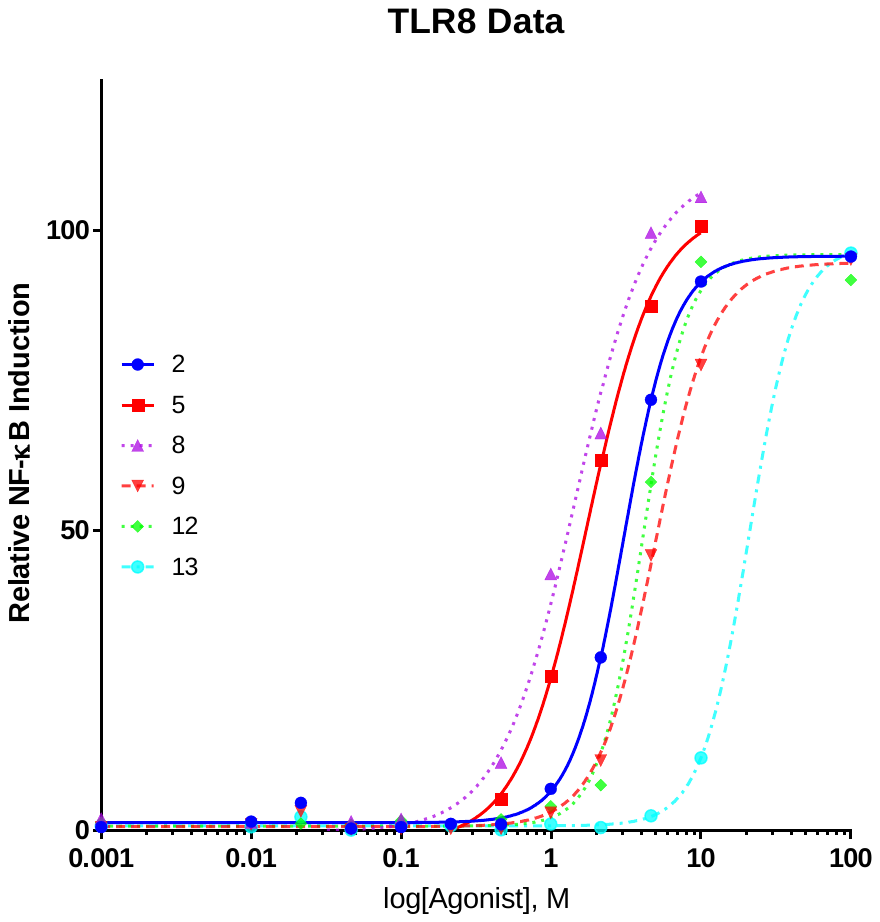

### **SI Figure 3. Cytokine Profile Data**

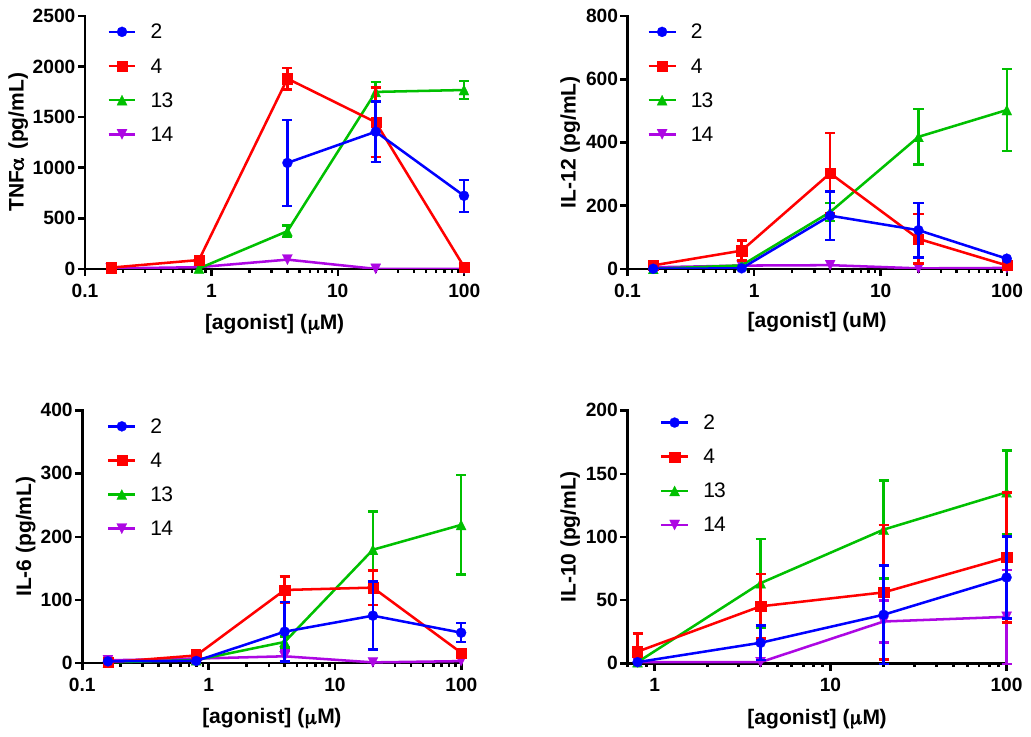

### **Compound Purity from HPLC Chromatographs.**

Chromatographs were acquired by UV detection (254 nm) with a Shimadzu LC2010 system using LCMSSolutions 3.4 (**4**, **6 – 14)** or EZStart 7.4 software (**5**). Compounds were eluted with 80:20:0.1 to 20:80:0.1 H_2_O:MeCN:HCOOH gradient at 0.6 mL/min for 12 minutes on a Supelco Discovery HS C18 (4.6x150mm, 3 micron) column at 30°C.

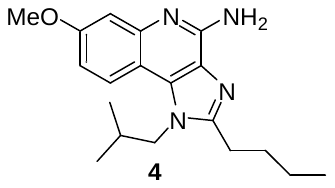

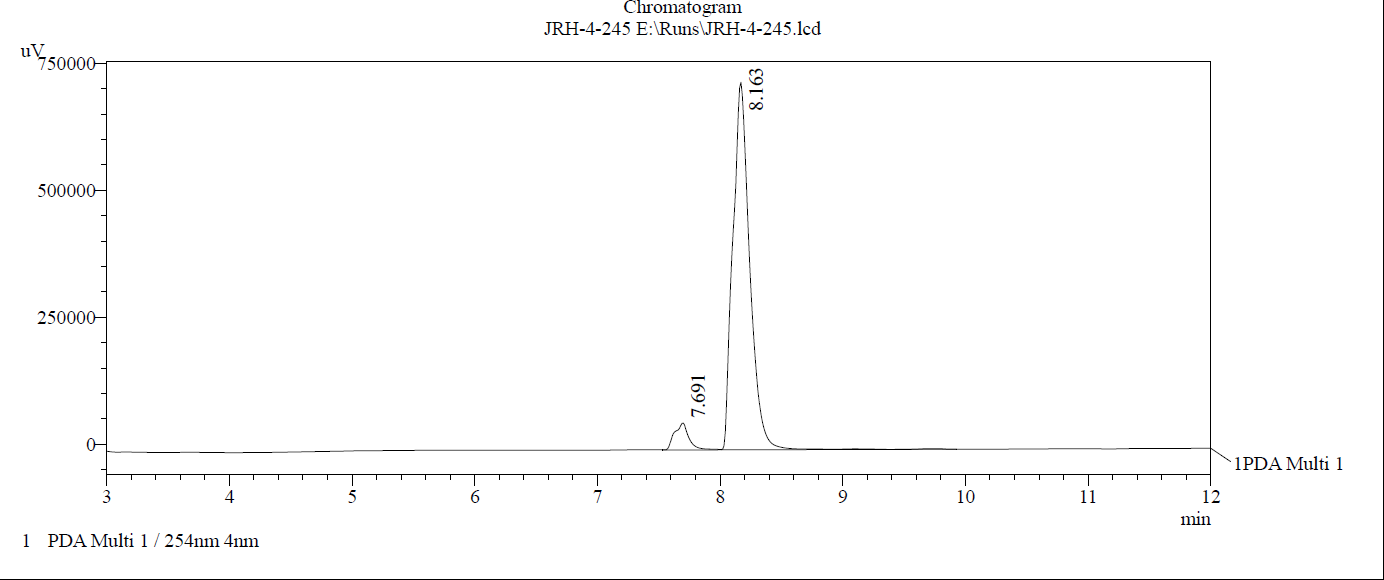

Purity: 94.1%

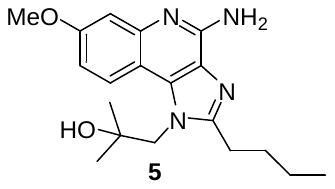

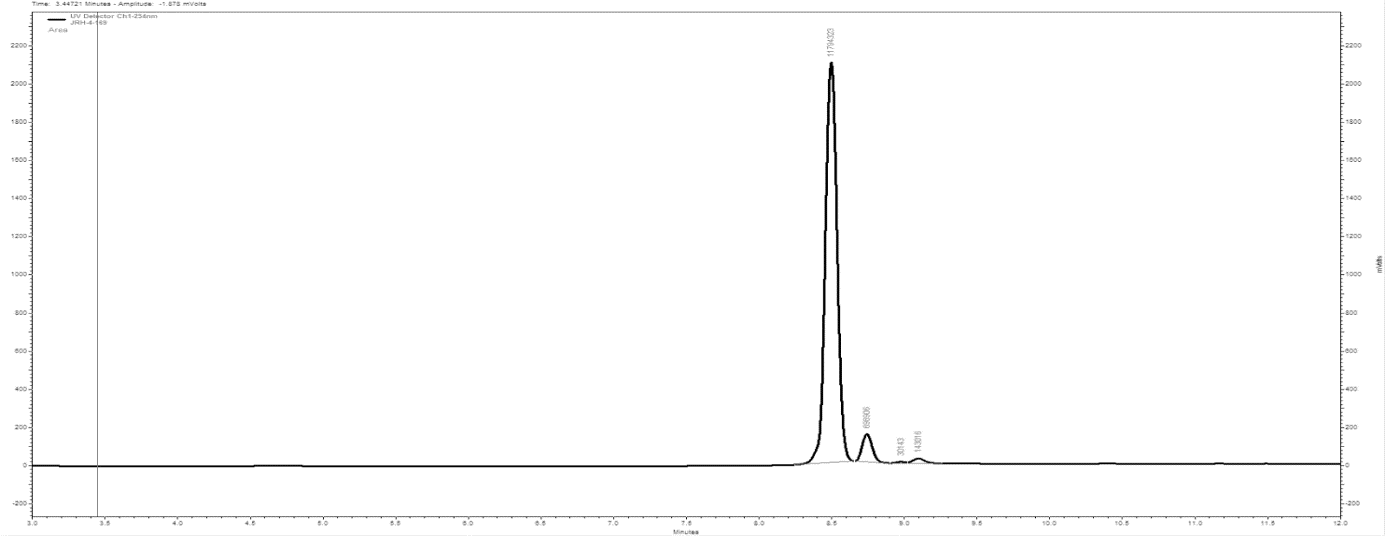

Purity: 93.5 %

9.08

8.71

8.48
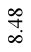

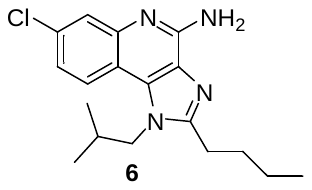

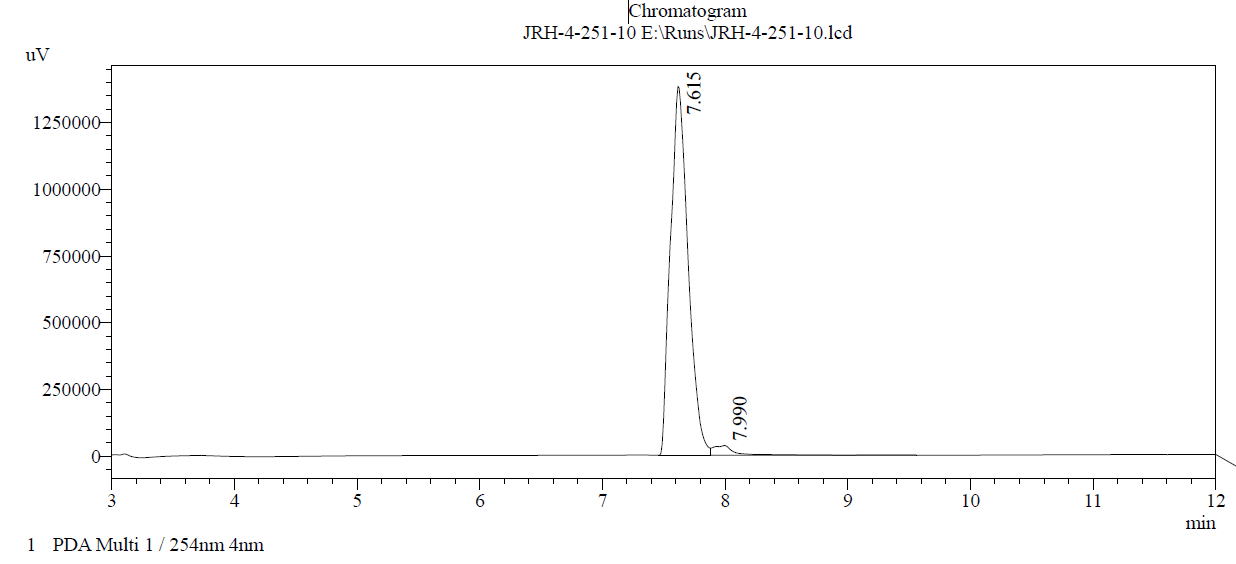

Purity: 98.2%

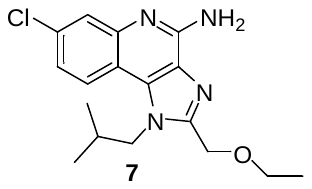

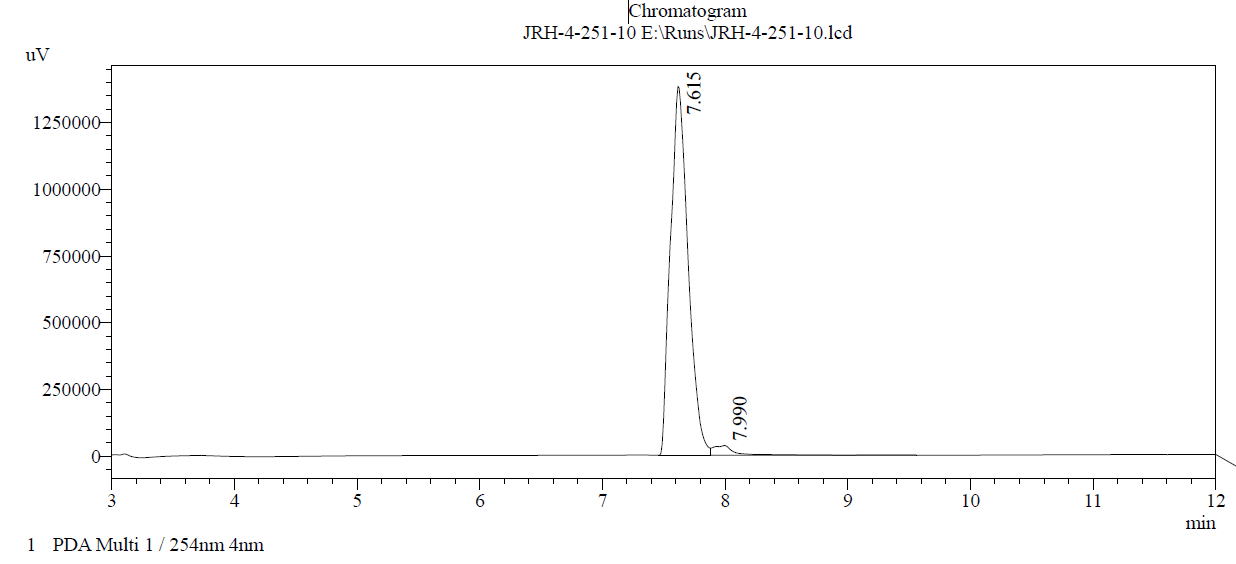

Purity: 96.9%

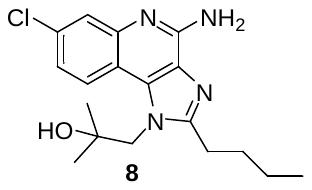

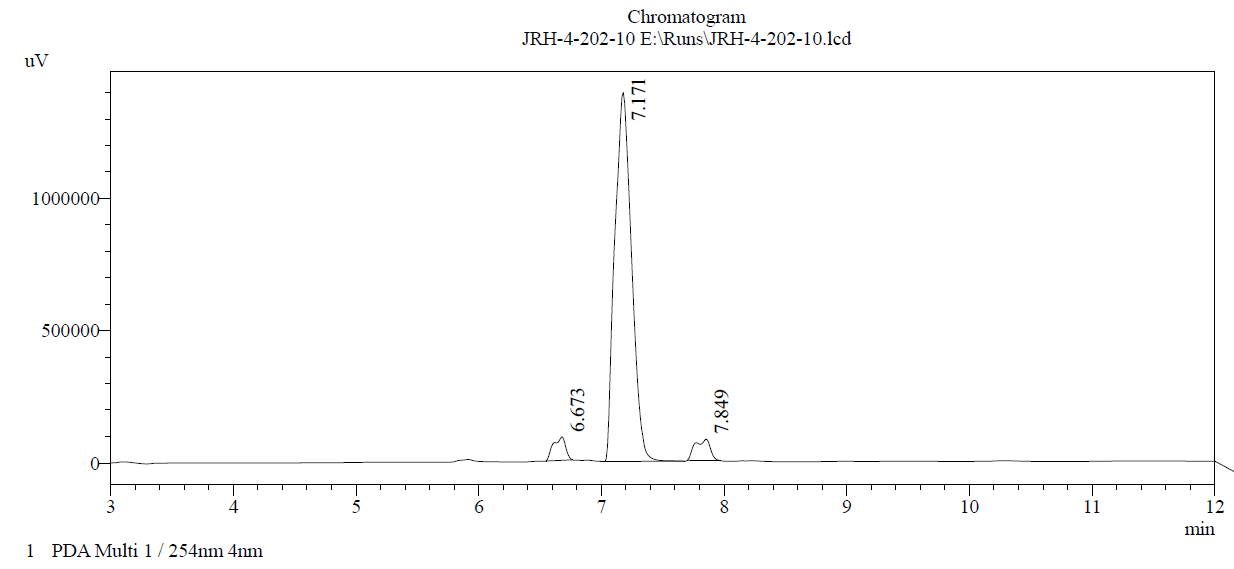

Purity: 91.0%

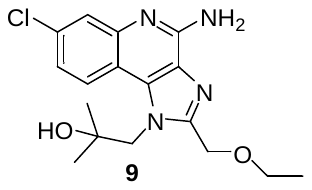

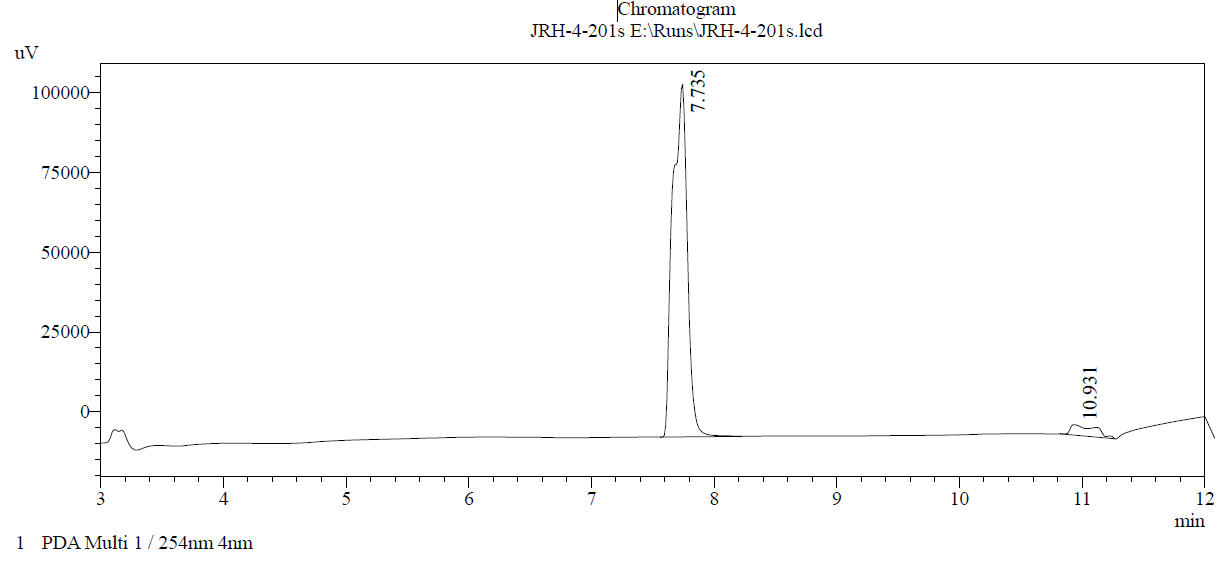

Purity: 95.0%

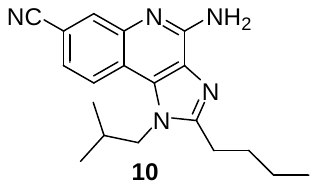

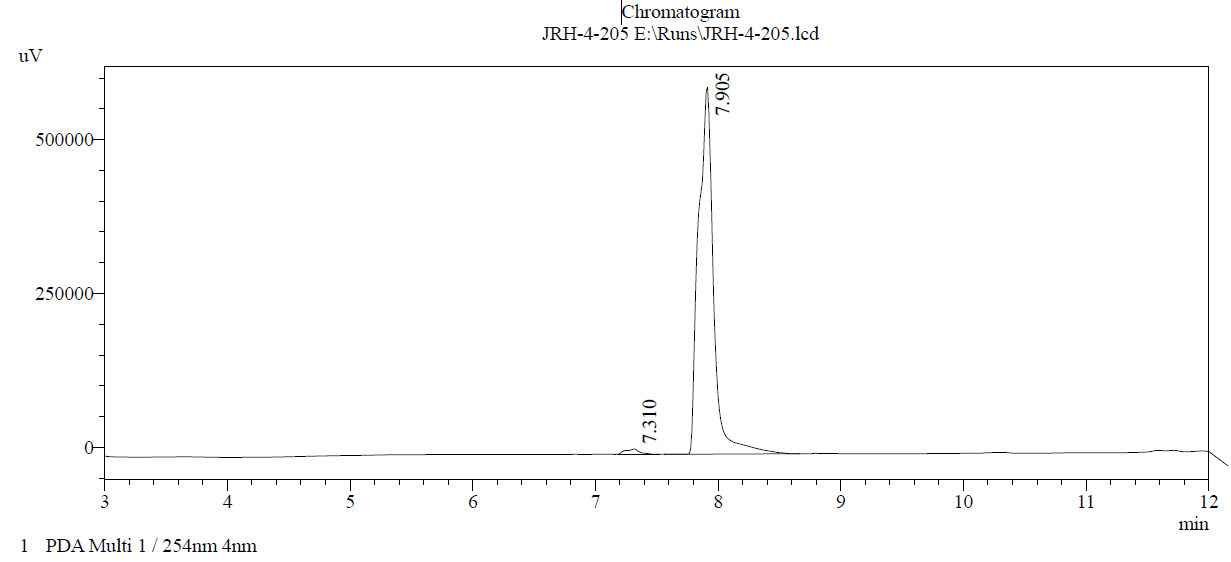

Purity: 98.6%

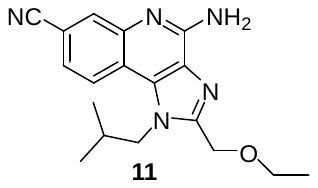

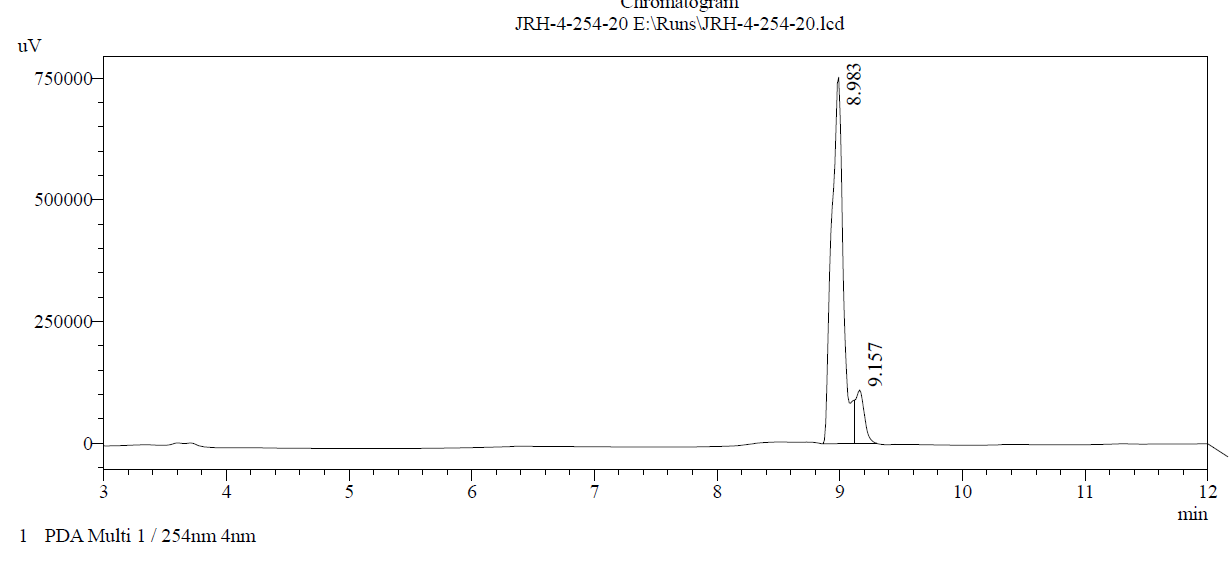

Purity: 90.4%

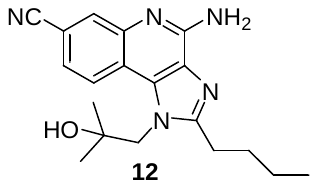

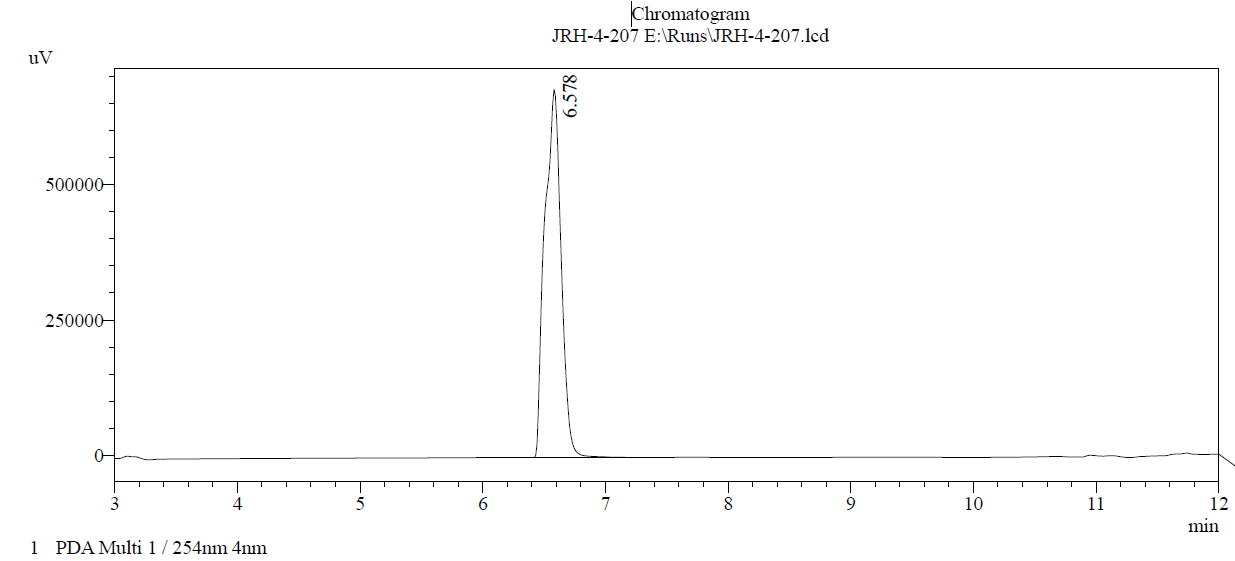

Purity: 99.9%

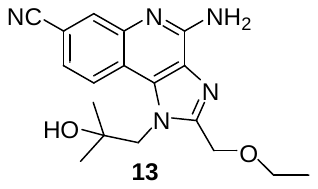

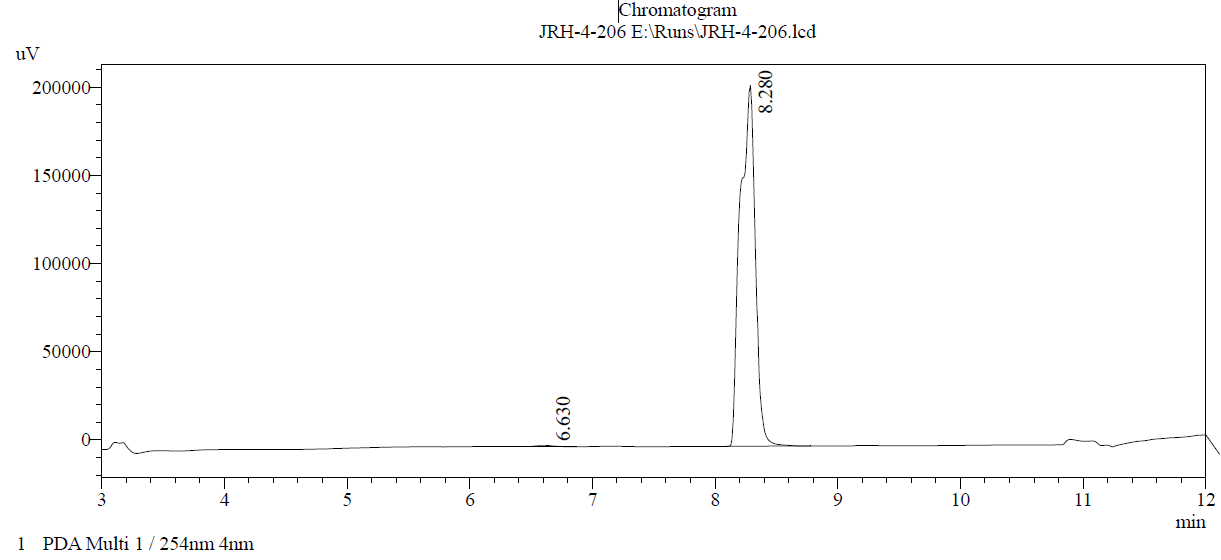

Purity: 99.7%

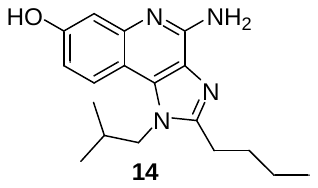

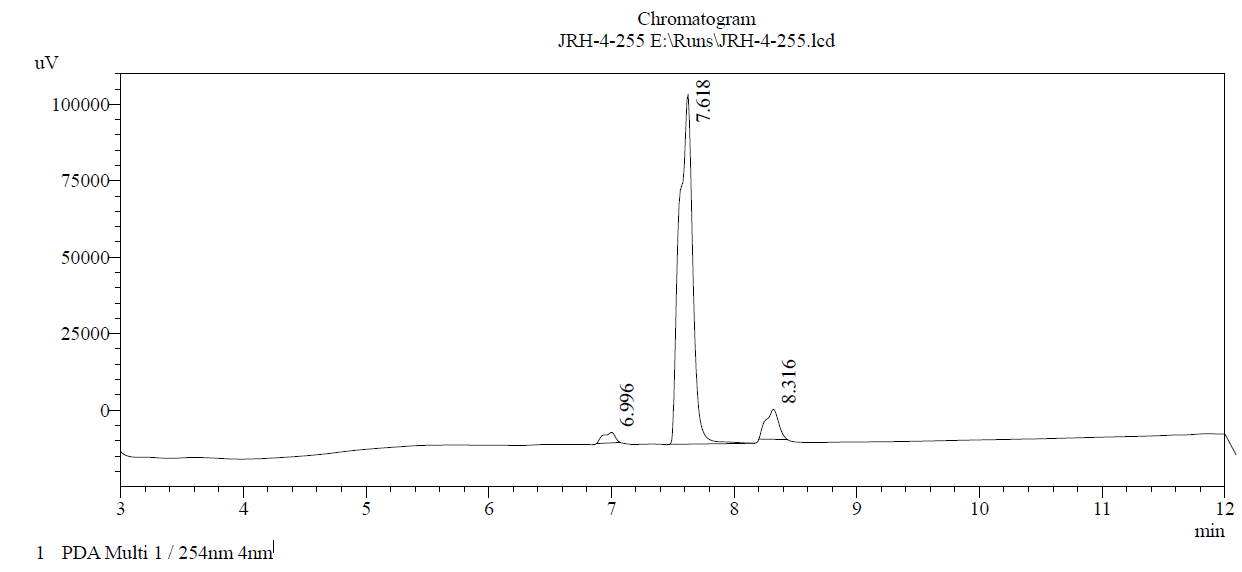

Purity: 90.2%
